# Optimizing microbial intake helps to maintain the gut microbiome diversity

**DOI:** 10.1101/2025.03.05.641598

**Authors:** Vitor M. Marquioni, Ann-Cathrin Hofacker, Jocksan V. Villavicencio, Florence Bansept

## Abstract

The animal gut holds a myriad of microbes with demonstrated importance for the host’s health. Properties of this microbiome have been studied in different host conditions in search of the best biomarkers for practical medical interventions. One well-studied quantity is the *α*-diversity, which usually correlates with lower values and pathological status. Diet plays an important part in the gut microbiome assembly. While its nutritional content has been extensively investigated, we ask instead how can the microbial content of food impact the gut *α*-diversity, taking into account the intermittent nature of feeding. We design a simple model that isolates the effect of intermittent microbial migration, for which we are able to develop an analytical treatment. Specifically, we find that there is a set of feeding parameters (feeding interval and food microbial content) that maximizes the time-averaged observed Shannon diversity, which we call the Optimal Feeding Strategy. Using a combination of numerical and analytical techniques, we show that, in the optimal scenario, diversity converges to that of the food, and that the optimal feeding rate converges to the average clearance rate. As probiotics become more and more widely used, we hope this work can help assess how quantitative ecological control can be used to improve intake protocols of live biotherapeutic products.

## 1 Introduction

The digestive system of animals is inhabited by many microbes [1, 2]. It is estimated that there are about 10^14^ bacteria in the human colon, a number comparable to the number of cells forming the human body [3]. These microorganisms are important to their hosts because they fulfill various functions. For instance, they participate in biochemical processes [4, 5], e.g. by digesting complex fibers [6], and take part in the immune function [7, 8], e.g. through competitive exclusion [9, 10]. An imbalance in the microbiome composition prevents it from performing these functions properly, and a wide range of diseases is associated to such imbalances [11–13], from metabolic disorders [14], like obesity [15, 16] and diabetes [17], to cancer [18] and psychological conditions [19, 20]. Measures of the gut microbiome *α*-diversity in different host’s conditions commonly show associations between poor health status and a less diverse microbiome [21]. Causal links between diversity and health remain to be generically understood; nonetheless, *α*-diversity can show some predictive power over patients’ outcome in some cases [22]. Therefore, it is of crucial interest to human health to get a better understanding of how the gut microbiome composition is regulated and its diversity sustained.

One important driver of the gut microbiome composition is the host’s diet [23–25]. Carbohydrates [26], fats [27], proteins [28], and other compounds, like alcohol [29] and food sweeteners [30] all have documented effects on the gut microbiome. Besides the abiotic content of ingested food, its microbial content [31, 32] has only been scarcely explored and yet could be of important medical relevance [33]. Some microbes present in the food indeed reach and settle in the gut [23, 32], where their interactions with the resident microbiome may have different outcomes. Food poisoning corresponds to cases when such interactions are harmful to the host [34], leading to adverse health symptoms. At the other end of this spectrum, the consumption of probiotics [5, 35] is an increasingly widespread therapeutic practice [36] aiming to improve individuals’ health, with promising results [37–39]. However, clear guidelines for a rationalized probiotics usage are still missing [40].

One common aspect of feeding throughout the animal kingdom is discontinuity, i.e., there are time intervals between meals [25], during which the amount of microbes that migrate to the gut is negligible – even though some microbes from the oral cavity might still reach it [41]. The impact of feeding temporal patterns on health has gained some attention over the last years, with an increasing number of shown associations between time-restricted feeding and important metabolic markers, like glucose and insulin blood levels [42, 43]. Noticeably, feeding time has been shown to be an important control of diurnal gut microbial oscillations in mice [44], although data with such high temporal resolution remain scarce, and little is known about microbiome fluctuations that result from feeding.

From a theoretical ecology perspective, migration has been shown to be an important mechanism to maintain the diversity of a focal community [45]. However, most models of microbial communities consider continuous migration terms [46, 47]. Here, we develop a model of gut microbial community where new microbes are introduced periodically in the system. We focus on the case where, in the gut, the different microbial types are equivalent competitors for a shared carrying capacity, which allows us to develop a full analytical framework. We derive the feeding parameters – feeding interval and food microbial content – that maximize the average Shannon diversity [48] of the community, which we call Optimal Feeding Strategy (OFS). We first focus on the conditions for the existence of an OFS by developing rigorous approximations, and we numerically show that it almost always exists when the number of types is moderately large (≳ 20). Then we look at the diversity achieved through the OFS and show it quickly converges to the food diversity as the number of microbial types increases. Finally, we show that the optimal feeding parameters also converge to analytically tractable values.

By isolating the effect of microbial migration through feeding, our work helps understand how it can be used as a control mechanism by the host to maintain its gut microbiome diversity, and associated health status. In particular, this paves the way to the development of controlled experiments aiming to optimize the protocols of live biotherapeutic treatments [49], like probiotics or capsulated fecal microbiota transplants [50].

## 2 Model and Methods

### 2.1 Model

We consider a model of a microbial community in which *S* microbial types compete for a shared carrying capacity *K*. Each type *i* has a maximum growth rate *r*_*i*_ and is cleared out of the system at rate *c*_*i*_. Writing the abundance of each type *n*_*i*_, the equations describing the system between feeding events write

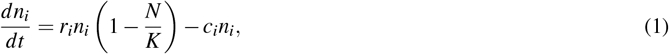

where *N* = ∑_*i*_ *n*_*i*_ is the total abundance of microbes. In the general case, the only stable non-trivial fixed point of this system of equations is given by *n*_*i*_ = 0 for all *i*≠*i*^∗^, and 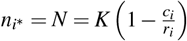 where *i*^∗^ is the microbial type with the smallest ratio 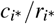 (see linear stability analysis in the supplementary material).

Feeding is modeled as instantaneous bursts of microbial immigration of size *n*_*f*_ every *τ* time intervals. Each microbe *i* is present in the food with a fixed fraction *f*_*i*_, such that *n*_*i*_(*t*) increases by *f*_*i*_*n*_*f*_ every *τ*, as shown in Figure 1. Feeding induces oscillations of the abundances with period *τ*. After a transient phase, the system stabilizes and abundances range from *n*_*i*_(*mτ*^+^) = *n*_*Li*_ + *f*_*i*_*n*_*f*_ to *n*_*i*_((*m* + 1)*τ*^−^) = *n*_*Li*_, with *m* = 1, 2, … and *n*_*Li*_ the basal level of oscillation of type *i*. We also define *N*_*L*_ = ∑_*i*_ *n*_*Li*_. In this regime, a direct integration of Eq.(1) over one period of oscillation leads to

**Figure 1.**
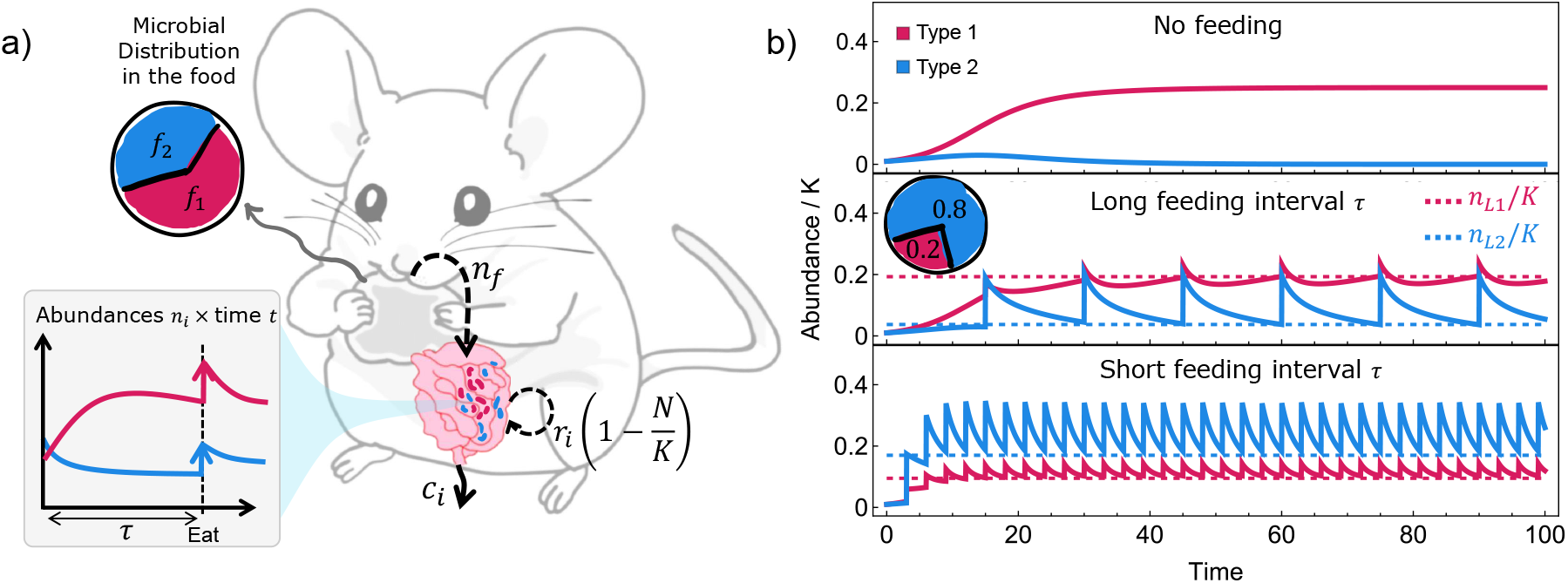
In an intermittent feeding model, feeding can rescue populations from extinction. In (a), we show an outline of the model in a two microbial types case. Within the gut, the abundances of the different types *n*_*i*_ change in time according to Eq.(1): they grow with maximal rate *r*_*i*_, share a carrying capacity *K* and are expelled from the gut at rate *c*_*i*_. At every time *τ, n*_*f*_ new microbes are introduced in the gut through feeding, changing the current abundances. A fraction *f*_1_ of *n*_*f*_ corresponds to type 1, while a fraction *f*_2_ = 1− *f*_1_ corresponds to type 2. In (b), the behavior of equation (1) in three different cases is shown: on top is the case in absence of feeding, where type 1 (pink) survives and type 2 (blue) gets extinct; in the middle, feeding rescues type 2 from extinction, with *n*_*f*_ = 0.2*K* and *τ* = 15; at the bottom, type 2 is not only rescued from extinction, but made more abundant than type 1 through feeding, with *n*_*f*_ = 0.2*K* but *τ* = 3. The dashed lines represent the basal levels of oscillations, calculated in Eq.(2). We used as relative abundances 20% for type 1 and 80% for type 2 in the food and *r*_1_ = *r*_2_ = 1.0, *c*_1_ = 0.75, *c*_2_ = 0.85, *K* = 1000.

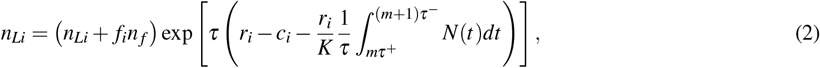

where we recognize the average of the total abundance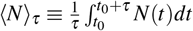. In what follows, we find a good approximation for ⟨*N*⟩_*τ*_, which allows us to analytically study our model.

Figure 1a shows an outline of the model, whereas Fig. 1b highlights the shifts in the microbial abundances when intermittent feeding is included. In the absence of feeding (upper plot), numerical integration of Eq.(1) shows type 1 surviving over type 2. The effective carrying capacity 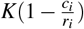 is smaller than *K* as a consequence of clearance. In the middle plot, feeding rescues type 2 from extinction, putting the system in an oscillatory state which is in phase with feeding. In the lower plot, by decreasing the feeding interval, type 2 can be made even more abundant than type 1.

### 2.2 Approximations for the average of the total abundance ⟨*N*⟩_*τ*_ (small *τn*_*f*_)

#### Trapezoidal approximation

We consider a trapezoidal approximation for the time-averaged abundances *n*_*i*_ over a period *τ*, given by ⟨*n*_*i*_⟩_*τ*_ ≈ *n*_*Li*_ + *f*_*i*_*n*_*f*_ */*2. This zero-th order approximation in *τ* is valid for 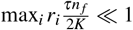 (see the supplementary material). From this, it results that in the long time limit,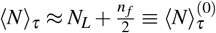. Isolating *n*_*Li*_ from equation (2), summing over *i* and replacing ⟨*N*⟩_*τ*_ by 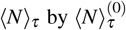, we find

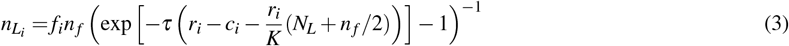

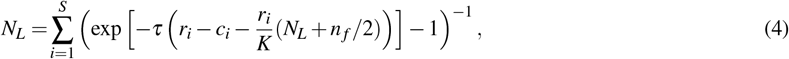

which can be solved numerically for *n*_*Li*_ and *N*_*L*_.

#### Continuous feeding limit (*τ* → 0 and *n*_*f*_ → 0)

Let us now suppose that *τ* → 0 and *n*_*f*_ → 0 while keeping *n*_*f*_ */τ* = *α* constant. From Eq.(4), we find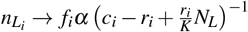. This limit coincides with the solution of the equation

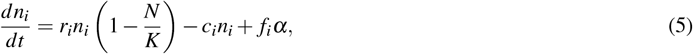

in which there is a continuous inflow of microbes with rate *α* to the microbiome, called *feeding rate* in the following. This result defines Eq.(5) as the *continuous feeding limit* of the model.

### 2.3 Diversity and Optimal Feeding Strategy (OFS)

We consider the time-averaged Shannon diversity 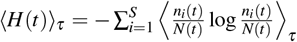 of the microbiome as a proxy for the host’s health. We aim to understand how microbial feeding impacts this diversity measure. In particular, we want to identify the set of parameters (*τ, n*_*f*_) that maximize it, which we call the *Optimal Feeding Strategy* (OFS).

In the regime where deviations around the averages ⟨*n*_*i*_⟩ are small (*n*_*f*_ and *τ* small), it is possible to show that ⟨*H*⟩_*τ*_ is well approximated by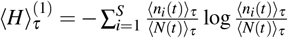, and it is exact in the continuous feeding limit, as *H*(*t*) converges to a constant value *H* in the long time limit. In addition, we also show that 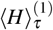 is a good approximation to 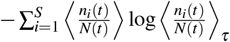 which is typically used empirically – as measuring relative abundances remains more common than obtaining absolute ones. A rigorous derivation of both results can be found in the supplementary material.

## 3 Results

Without feeding, in the long time limit, the Shannon diversity of the community is equal to 0. With feeding, in the trapezoidal approximation regime (see section 2.2), where deviations around the average abundances are small, we can write

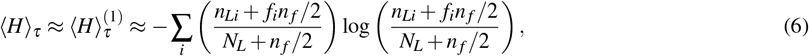

which is greater than 0, even when all but one *n*_*Li*_ is close to zero.

### 3.1 Feeding can lead to the existence of an OFS

Let us consider a two-microbial types case in which type 2 gets extinct in the absence of feeding. As depicted in Fig.1b, the periodic arrival of microbes can rescue it from extinction, increasing its average abundance over a time window *τ*. However, if the feeding interval *τ* is small, the number of microbes that reach the gut from feeding becomes so important that it takes over local interactions. That can result in a lower diversity, if, as in Fig.1b, type 1 is rare in the food. Thus, in this case, an intermediate feeding interval is bound to maximise the time-averaged diversity: feeding needs to be frequent enough to rescue type 2, but not too frequent that we loose type 1.

In Figure 2a, the heatmap shows the diversity ⟨*H*⟩_*τ*_ numerically obtained for different values of *τ* and *n*_*f*_ for an *S* = 2 case. The curve in white shows the set of points where diversity is maximal, i.e., it is the OFS. Analytical results on this curve are discussed in the following sections, but as a first approach, we can numerically solve equations (3) and (4) and use the results in Eq.(6). This procedure gives the black curves of Figure 2b, which are in good agreement with the numerical evaluation of the model for small *τn*_*f*_. In the rest of the paper, we investigate how general is the existence of an OFS.

**Figure 2.**
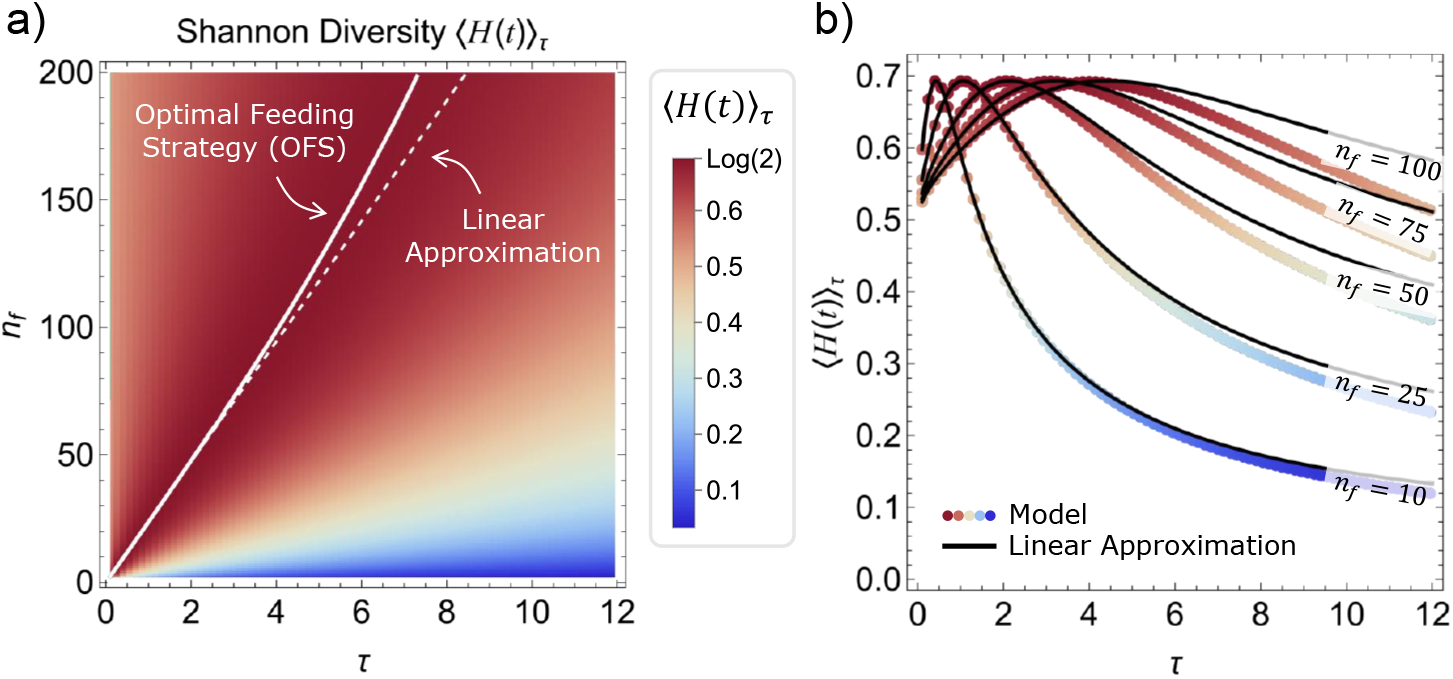
An Optimal Feeding Strategy exists and can be well approximated in a 2 microbial types case. In (a), we numerically calculated the time-averaged Shannon diversity ⟨*H*(*t*) ⟩ _*τ*_ for long simulation times, which is plotted as a heatmap. For each *τ*, we identified the value *n*_*f*_ for which ⟨*H*⟩ _*τ*_ was maximal, thus defining the Optimal Feeding Strategy, shown by the white curve. The dashed curve shows the linear approximation obtained from Eq.(7). In (b), we plot ⟨*H*⟩ _*τ*_ as a function of *τ* for different *n*_*f*_, i.e., single lines of the heatmap in (a). The black curves are given by Eq.(6) where *n*_*Li*_ are given by the trapezoidal approximation. Simulation parameters are: *r*_1_ = *r*_2_ = 1.0, *c*_1_ = 0.75, *c*_2_ = 0.85, *K* = 1000 and *f*_1_ = 0.2.

### 3.2 The OFS admits a linear approximation

An OFS can also be searched for in the continuous feeding limit. In this case, the *linear OFS approximation* is defined by the *n*_*f*_ = *ατ* line with the value of *α* that maximizes the Shannon diversity *H* – which is, in this case, constant in the long time limit. Notice that the total abundance *N* depends on *α*, and thus a *feasible* linear OFS requires that both *α* and *N* are positive quantities.

#### The special case

*S* = 2 When *S* = 2, we can use that *n*_1_ = *n*_2_ = *N/*2 maximizes the Shannon diversity, and applying it on the fixed point of equation (5), we find

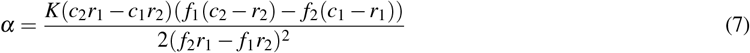

as the optimal feeding rate, with the total abundance given by

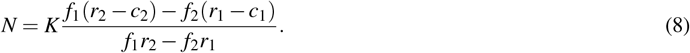

In Fig. 1, the dashed white line shows the linear OFS calculated with this equation. A joint analysis of these equations gives the following conditions for feasibility:

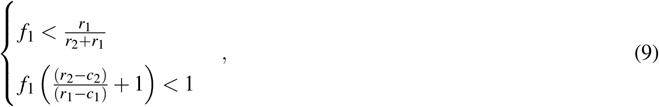

in which we considered *r*_1_*/c*_1_ *> r*_2_*/c*_2_ without loss of generality. Notice that these conditions set an upper bound to the fraction *f*_1_ of type 1 in the food. If *f*_1_ is greater than this bound, feeding cannot be used as an effective strategy to maximize diversity.

#### The general solution of the linear OFS

When the number of microbial types *S* is greater than 2, the diversity can also be optimized in the continuous feeding limit, but, in general, not to log *S*, which is the maximal Shannon diversity for a community with *S* types. This is because the number of equalities *n*_1_ = *n*_2_ = … = *n*_*S*_ to be satisfied to reach *logS* is larger than the number of variables in the system, which are, in the framework of the continuous limit, *α* and *N*. Hence we use a Lagrange multiplier approach to the following problem: the maximization of

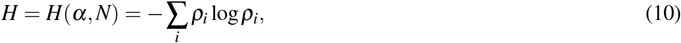

where 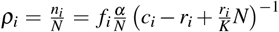 is the fixed point of Eq.(5), and subjected to the constraint ∑_*i*_ *ρ*_*i*_ = 1. Using the Lagrangian ℒ (*α, N, λ*) = *H*(*α, N*) − *λ* (∑_*i*_ *ρ*_*i*_ − 1), the equation 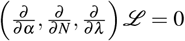 gives the optimal solution for *α*,

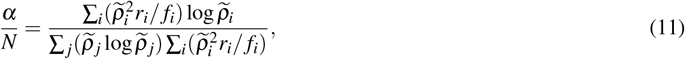

where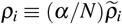, and the maximal diversity is given by

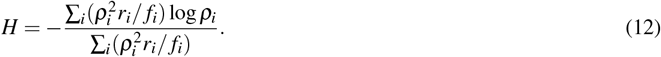

Equation (11) can be solved numerically with the constraint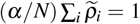, and we provide an algorithm for that in supplementary material.

#### The existence of a linear approximation conditions the existence of an OFS

In the long time limit, the total abundance *N*(*t*), between two consecutive feeding events, is a decreasing convex function of time. As a result, the existence of a feasible linear OFS is a sufficient condition for the existence of a feasible OFS in the presence of intermittent feeding, i.e., if the linear OFS is feasible, there is a curve (*τ, n*_*f*_) that maximizes ⟨*H*⟩_*τ*_. We show both results in the supplementary material. As a consequence, the conditions (9) are also valid when feeding is intermittent.

### 3.3 Numerically solving the linear OFS approximation

We investigated the numerical solutions of Eq.(11) using randomly generated communities with the richness ranging from *S* = 2 to *S* = 1000. For a given *S*, we drew the parameters *r*_*i*_, *c*_*i*_ and *f*_*i*_ at random and numerically solved Eq.(11) for *α*. We considered growth rates *r*_*i*_ drawn from the log-normal distribution [51] *log𝒩* (*µ* ≈ −1.5, *σ* ≈ 1.4) – giving growth rates expressed in per hours – which adjusts the data from [52] for microbial generation times at optimal growth temperature (*r* = log 2*/*(Generation Time)). For the clearance rates, we first considered *c*_*i*_ = 0.02*/*hour ≈ 0.5*/*day for every *i*, i.e. half of the microbial content in the gut is expelled per day [53], but wider distributions were also studied. Each *f*_*i*_ was first uniformly drawn from the interval [0,1], and then normalized to 1 (resulting in a Beta distribution 170 (Beta(1,*S*™1)) for each f_*i*_). Variations around these distributions were also considered, but without significant changes on the results, apart from particular cases discussed in Section 3.5 (see the supplementary material for a more systematic exploration of these distributions). Notice that Eq.(11) and the constraint 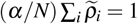 are equations on the variables *α*/*N* and *N*/*K*; therefore, the carrying capacity does not need to be defined to find the linear OFS.

For each richness *S*, we drew 420 random communities and computed the optimal solution of *α*/*N* for each of them. The rare cases with numerical convergence problems were discarded (see supplementary Fig. S2) and we computed the proportion of the solutions that were feasible (see supplementary Fig. S3). Fig. 3 thus summarizes three important results:

**Figure 3.**
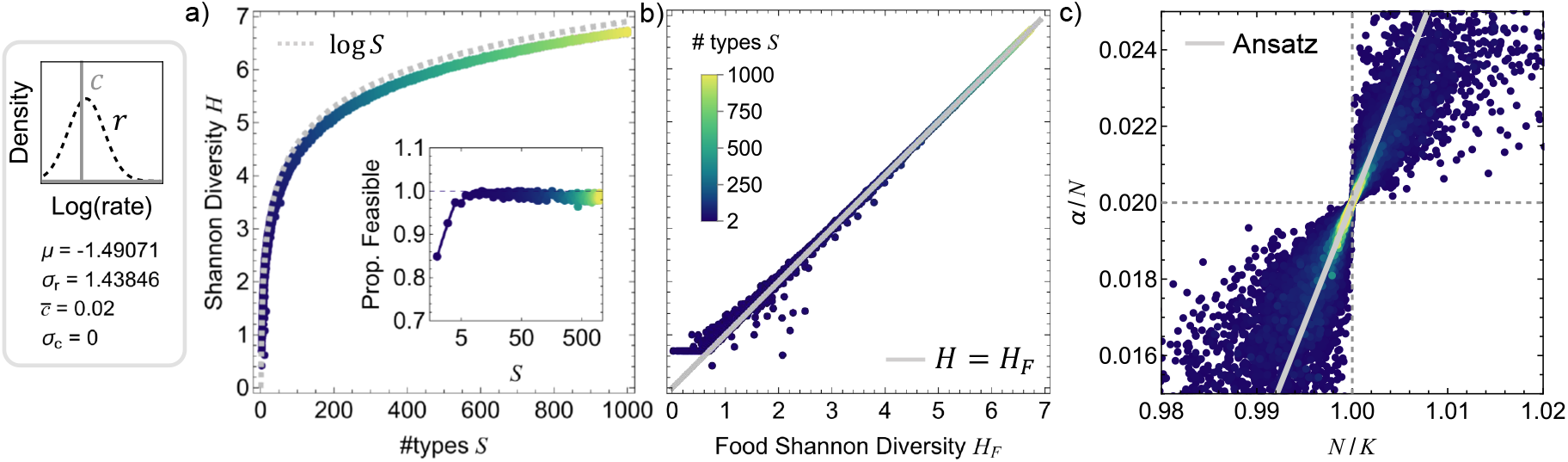
Optimal parameters converge to the ansatz solution in the large S limit. The result of the diversity maximization problem are shown for randomly generated communities (with *r*_*i*_ ∼ *log* 𝒩 (μ,σ_r_) and 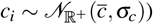 for an increasing number of microbial types *S* (shown with the color scale). In (a), the inner plot shows the proportion of random communities that display a feasible Optimal Feeding Strategy that approaches 1 as *S* increases, and the outer plot shows the maximal diversity *H* that is actually achieved by these communities. In (b), the maximal achieved H as a function of food diversity *H*_*F*_ is shown to approach *H*_*F*_ as *S* increases. In (c), we plot all the parameter pairs (*N/K,α/N*) that maximize diversity when a feasible OFS exists. As the number of types increases, the solutions accumulate around the gray line (given by Eq.(14)) and converge to *N* = *K* and *α* = ⟨*c*_*i*_⟩*N*. Parameters for the simulations are shown in the left panel.

#### The proportion of feasible maximal solutions increases for large numbers of microbial types

As the richness *S* increases, the proportion of feasible solutions increases fast to 1, and this result is observed for a large range of parameters (see supplementary Fig. S5), showing that the existence of an optimal solution becomes almost certain for large microbial pools, supporting the hypothesis of microbial feeding as a way of diversity control in microbial communities.

#### The maximal diversity correlates to the diversity in the food

The correlation between the maximal diversity in the gut and the diversity in the food is another clear result. Whereas in the *S* = 2 case, the diversity *H* can be maximized to higher values than what is observed in the food *H*_*F*_, the numerical results point to a “saturation effect” when *S* is large, that limits the maximal diversity to *H*_*F*_. This result is consistent with further theoretical results (see Section 3.4) and was observed for a large range of parameters (see supplementary Fig. S7).

#### The optimal feeding rate converges to the clearance rate for large numbers of microbial types

As the number of types increases, the optimal feeding rate *α* converges to ⟨*c*_*i*_⟩*N* and *N* to *K*, with decreasing deviations from this point

### 3.4 An ansatz to the linear OFS approximation

In light of the numerical results shown in Fig. 3, we propose an ansatz to the optimal diversity for large *S*, writing 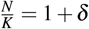, and assuming *c*_*i*_ = *c* for every *i*. We expand the Shannon diversity *H* = −∑_*i*_ *ρ*_*i*_ log *ρ*_*i*_ in *δ*, with *ρ*_*i*_ given by Eq.(10), around *δ* = 0, and we are able to show that, for uncorrelated distributions of *f*_*i*_ and *r*_*i*_,

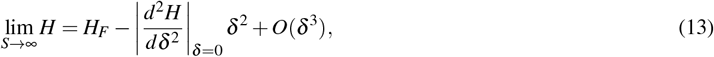

which means that *H* has a maximum at *δ* = 0, which equals to *H*_*F*_ (the detailed calculations are in the supplementary material). Moreover, for *δ* ≪ 1, from the constraint ∑_*i*_ *ρ*_*i*_ = 1, we find that

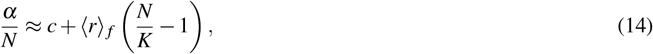

and thus *α/N* → *c* in the considered limit. The line defined by Eq.(14) is plotted as the continuous gray line in Fig. 3c.

### 3.5 Different growth and clearance rate distributions affect the convergence to the ansatz solution

Figure 4 shows how the growth and clearance rate distributions affect the obtained results. In Fig. 4a, changing the clearance rate distribution so that the *c*_*i*_ values are typically smaller than the *r*_*i*_ values can decrease the feasibility of the maximal solution, i.e. the proportion of cases for which feeding can maximize the Shannon diversity. In Fig. 4b, a broader clearance distribution significantly slows down the convergence to the ansatz solution when *S* increases, so that convergence to the grey line of Eq.(14) is visible only around the (*N/K, α/N*) = (⟨*c*_*i*_⟩, 1) point. In addition, Fig. 4c shows that, for growth rates much larger than the clearance rates, the solutions accumulates around the line given by equation (14), but without a clear convergence to the point (*N/K, α/N*) = (⟨*c*_*i*_⟩, 1).

**Figure 4.**
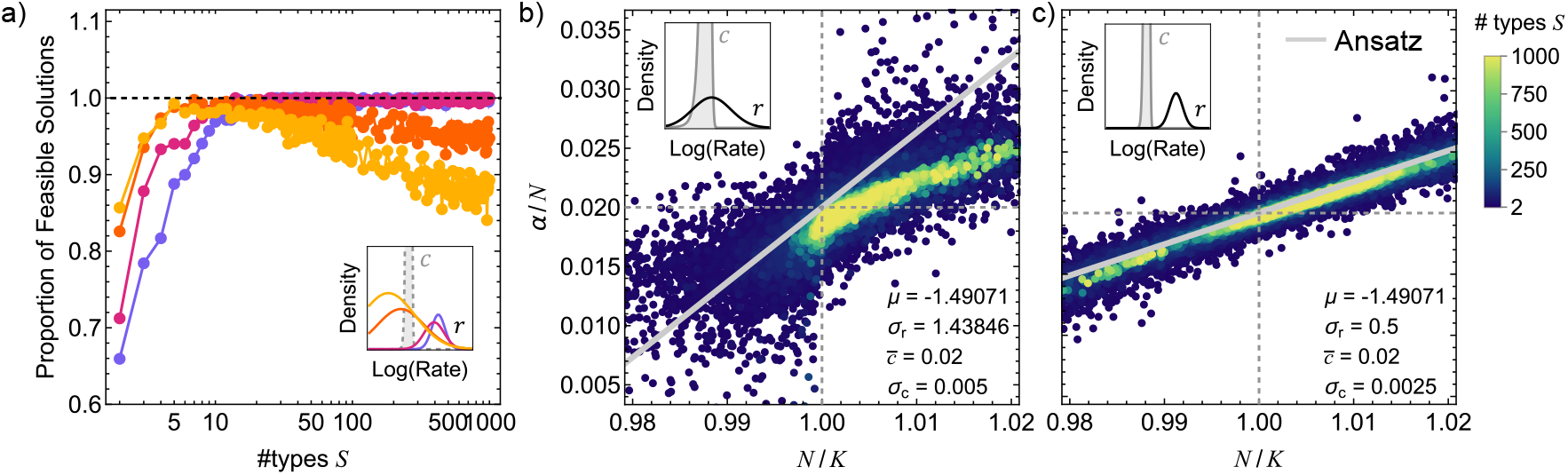
Clearance and growth rate distributions affect the feasibility and the convergence to the ansatz solution. In (a), we show how the proportion of feasible solutions is affected when the growth rate distribution changes while the clearance distribution remains unchanged. Feasibility decreases for large S when the growth rate becomes typically smaller than the clearance rate. Parameters used are 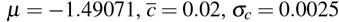 and *σ*_*r*_ = 0.5 (purple), 0.75 (magenta), 1.75 (orange) and 2 (yellow). In (b), a broader clearance distribution than in Figure 3c slows down the convergence to the ansatz solution and in (c), where the growth rates are significantly higher than the clearance rates, the solutions get closer to the gray line given by Eq.(14) but (*α/N,N/K*) do not converge to (1, ⟨*c*_*i*_⟩) as fast as when σc = 0.

We understand these points better by considering *c*_*i*_ ≡ ⟨*c*_*i*_⟩ + Δ*c*_*i*_ and further developing the ansatz solution, which gives

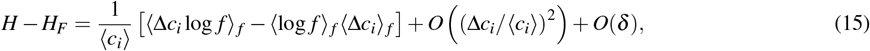

where the term in square brackets tends to zero for *S* → ∞ (detailed calculations can be found in the supplementary material). This shows that, for a finite number of microbial types, the zero-th order term in *δ* is indeed affected by the clearance distribution.

## 4 Discussion

In this work, we wanted to understand whether hosts can use feeding as a control mechanism [54] to favor the maintenance of diversity in their gut microbiome, which has been shown to be associated to health status [5, 21]. More specifically, we developed a mathematical model to investigate how the intermittent immigration of microbes affects the *α*-diversity of a community where different microbial types compete for a shared carrying capacity. We have shown that migration can rescue microbial types from extinction and that, in most cases, there even exists an Optimal Feeding Strategy (OFS), characterized by a relationship between feeding interval and food microbial content, that maximizes the average of the microbial diversity over time. We developed a linear approximation to the OFS and showed that the existence of this approximation is a sufficient condition for the existence of an OFS.We showed that an OFS almost always exists as soon as the number of types is moderately large (≳ 20), and that the diversity that is achieved through it quickly converges to the food diversity. In addition, we showed that the optimal feeding parameters also converge to values that we were able to explain analytically, although convergence may be slowed down by widening the distribution of clearance rates. Importantly, we have shown that the diversity reached by the system only mildly depends on the feeding temporal pattern, as it is almost entirely determined by the feeding rate, defined as the microbial content of food reaching the gut per unit of time.

Of course, an important limitation of our approach is not taking into account the nutritional content of food [25] explicitly: we choose a model where nutrients’ limitation is only accounted for through an effective interaction term that limits microbial growth when abundances reach a shared carrying capacity. This modeling choice presents two shortcomings: first, it assumes that all microbial types share a unique ecological niche, while, in reality, different nutrients’ sources can help sustain diversity in a way that is not taken into account here [55]. However, our model could alternatively also describe the optimization of live biotherapeutic products intake, like probiotics and capsulized fecal material used in fecal microbiota transplantation, in which there are not additional nutritional content. Second, we expect the amount of nutrients available to microbes to be strongly impacted by the host, since most nutrients will be absorbed through the gastrointestinal tract, creating a strong longitudinal gradient of nutrients, microbial, and pH concentrations in the gut [56–58]. Our model, on the other hand, assumes a well-mixed, chemostat-like system for simplicity, which could be a better description of specific anatomical compartments, like the cecum, which, in some animals that ferment cellulose, is somewhat isolated from the longitudinal flow [59, 60].

In addition to the conceptual limitations of our model, it is interesting to question whether the orders of magnitude for the optimal feeding parameters we found are realistic. Humans produce around 30g/day of dry feces, in which there are around 4 × 10^11^ bacteria/g [61, 62], resulting in ⟨*c*⟩*N* ∼ 10^13^ bacteria expelled per day. Therefore, in the context of our model, a human would need to ingest 10^13^ bacteria everyday in order to maximize its gut diversity. The estimated values for daily intake are rather in the range of 10^6^ to 10^9^ for a US diet [31], and only a fraction of this is expected to reach the gut, as the stomach acidity [63] and duodenum bile [64] create a barrier that kills many microbes. Even highly concentrated probiotics formulations have daily doses smaller than 10^12^ CFU [65]. However, even if our model was able to capture all the important mechanisms at play with diversity control in the gut, it does not necessarily mean that we would expect diversity to be optimized in all animal hosts – which could explain the discrepancy with our results. If indeed diversity is not optimized, then it becomes interesting to investigate why, from an evolutionary perspective. Controlled data on other animal hosts’ feeding habits is much scarcer, but we expect microbial ingestion to be proportionately higher for necrophagic animals or those with coprophagic behavior [66, 67], for which microbial feeding could have a more significant effect on gut diversity, in the light of our results. More controlled experiments would be required to test our predictions empirically, varying the microbial content of food and feeding patterns while maintaining nutrient intake constant, ideally with a wide range of host organisms. A good animal host candidate to start with may be fruit flies [68], in which microbiome complexity is low, and for which feeding assays have already been developed [69, 70].

In our reductionist approach, we used diversity averaged over time as a single indicator of the gut microbiome status. Although there are good empirical evidence that lower diversity is associated to worse health status, the causal nature of this association is not well understood yet [21]. To go beyond this approach, a functional description needs to be developed, which is challenging, although promising work goes in this direction, both experimentally and theoretically [4, 71]. An intermediate step in this direction could be to extend our model to a Generalized Lotka-Volterra (GLV) framework [47, 72, 73], in which one can take into account different types of (positive or negative) interactions between microbes. Indeed, in our model, diversity can only be sustained through migration, which is known as a mass effect by theoretical ecologists [74]. On the other hand, GLV would allow us to explore different regimes, in which multi-types equilibria exist even in the absence of feeding, and where feeding could make the system switch from one equilibrium to another. Another complex out-of-equilibrium dynamics observed in different GLV regimes with migration is the induction of chaotic species turnover [46], which is observed in *in vitro* experiments [75] and might become relevant for gut microbiome research when better time-resolved data become available.

Ecological control has been highlighted before as a possible microbiome edition tool [76]. In this work, we showed how, indeed, a global property of the community (the Shannon diversity) can be optimized by parameters that are under active control of the host. Although progress remains to be made in understanding what characterizes a healthy microbiome [77, 78], we believe that our model opens new perspectives for improving the use of ecological interactions as a control strategy of the gut microbiome [10, 54], in line with the personalized nutrition concept [25, 79].

## 5 Acknowledgments

We thank members of the M3g group for insightful discussions, especially Florian Labourel. The project leading to this publication has received funding from France 2030, the French Government program managed by the French National Research Agency (ANR-16-CONV-0001) and from Excellence Initiative of Aix-Marseille University - A*MIDEX. FB thanks Román Zapién-Campos, Ana Teles and Thomas Roeder for discussions that led to the onset of the project.

## 6 Author contributions statement

FB conceived the project, A-CH and JVV worked on an earlier version of the model. VMM extended the original model, performed the numerical and analytical study and analyzed the results. VMM wrote the original draft, VMM and FB edited it. All authors approved the final version of the manuscript.

## 7 Code availability

The codes used in this work are available at https://github.com/M-3-Group/IntermittentFeeding.

