## Supplementary material for "Optimizing microbial intake helps to maintain the gut microbiome diversity"

#### Contents

S1 Supplementary Images 2

S2 Linear Stability Analysis of the abundance equations in the absence of feeding 9

S3 The convexity of the total abundance: an important lemma 10

S4 The error of the trapezoidal approximation to the average total abundance 10

S5 Showing that a feasible linear OFS is a sufficient condition to the existence of a feasible OFS 11

S6 Approximations of the time-averaged Shannon diversity 12

S7 Maximizing the Shannon diversity in the continuous feeding limit for $S > 2$ 14

S8 The numerical method to solve the linear OFS approximation equations 16

S9 An ansatz solution to the linear OFS approximation equations 17

S10 The ansatz when the clearance rates are broadly distributed 20

### S1 Supplementary Images

(a) $r_i \sim LogNormal(\mu, \sigma_r)$

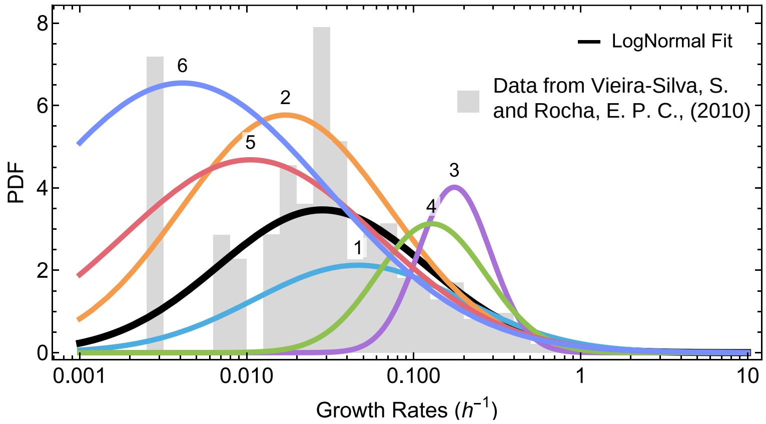


| Curve | $\mu$ | $\sigma_r$ |
|---|---|---|
| Fit | -1.49071 | 1.43846 |
| 1 | -1 | 1.43846 |
| 2 | -2 | 1.43846 |
| 3 | -1.49071 | 0.5 |
| 4 | -1.49071 | 0.75 |
| 5 | -1.49071 | 1.75 |
| 6 | -1.49071 | 2 |

(b) $c_i \sim Normal_{\mathbb{R}^+}(\bar{c}, \sigma_c)$

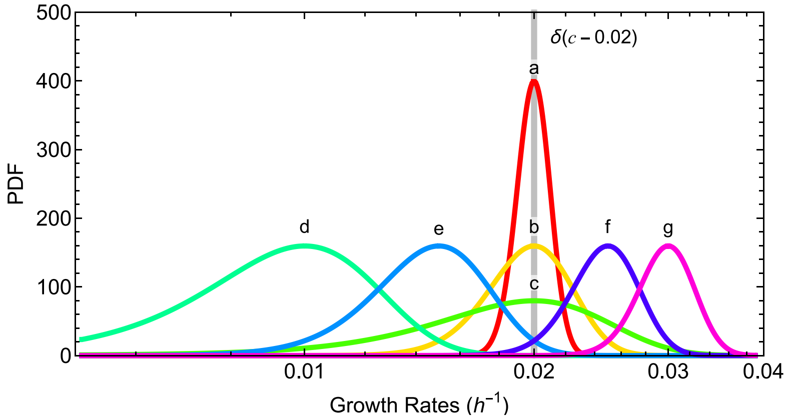


| Curve | $\bar{c}$ | $\sigma_c$ |
|---|---|---|
| $\delta$ | 0.02 | 0 |
| a | 0.02 | 0.001 |
| b | 0.02 | 0.0025 |
| c | 0.02 | 0.005 |
| d | 0.01 | 0.0025 |
| e | 0.015 | 0.0025 |
| f | 0.025 | 0.0025 |
| g | 0.03 | 0.0025 |

**Figure S1. Clearance and growth rate distributions used for numerical investigations.** In this figure, we show the different growth (a) and clearance (b) rate distributions used to numerically investigate the solutions of Eq.(3) from the main text. For the growth rates $r_i$, we considered log-normal distributions with parameters $\mu$ and $\sigma_r$, and for the clearance rates $c_i$, truncated normal distributions with original parameters $\bar{c}$ and $\sigma_c$. The histogram in (a) shows the data from [1], which agglomerates growth rates at optimal temperature of many microbial species from different environments, over which we adjusted a log-normal curve. In the following supplementary figures, each panel represents the numerical results considering one distribution from (a) and one distribution from (b).

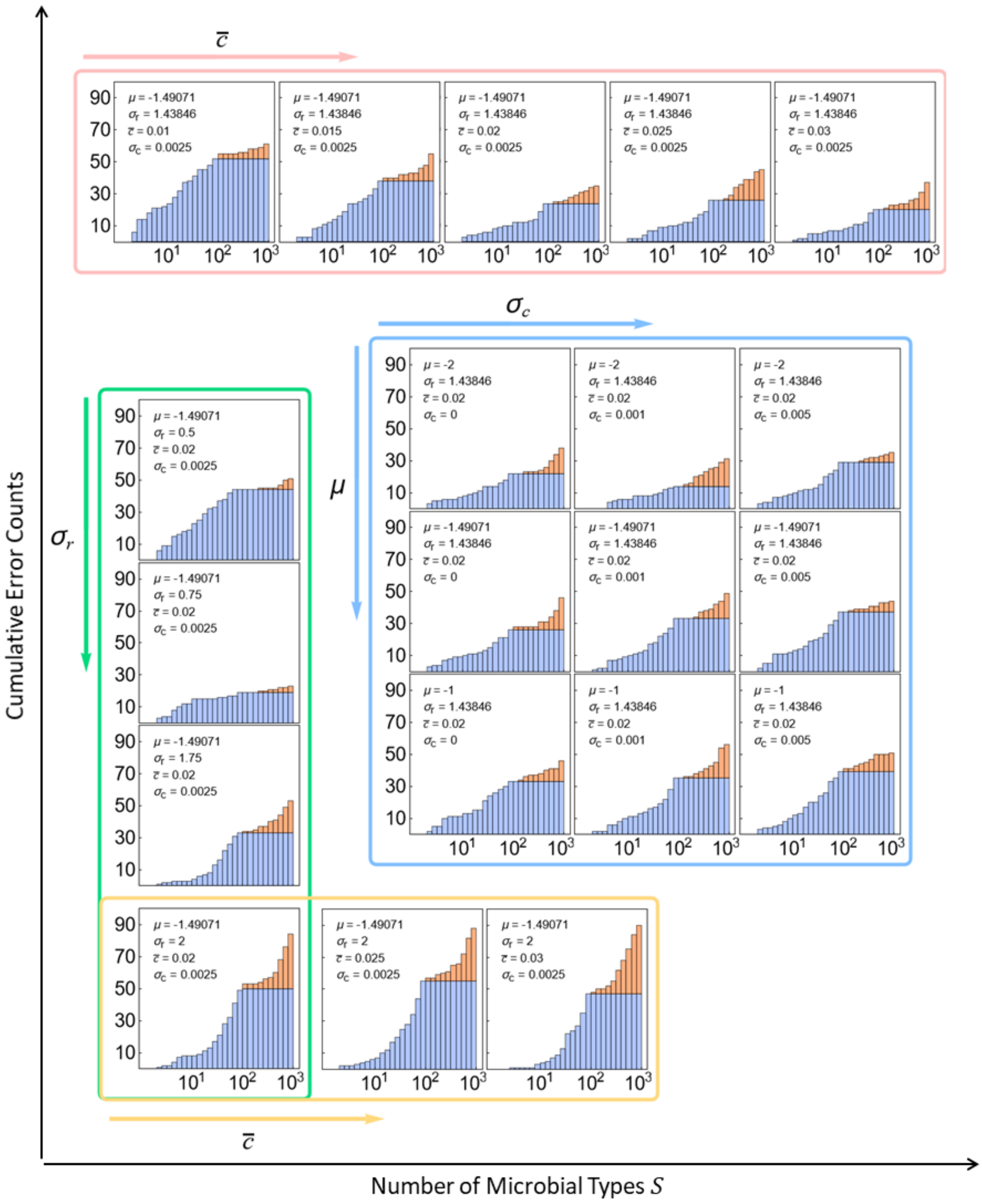


**Figure S2. The numerical method generates a number of errors that is negligible compared to the number of converging solutions.** In each panel, the cumulative number of errors (convergence problems due to numerical precision, see Section S8) that occurred while numerically solving Eq.(11) from the main text is shown as a function of the number of types $S$. We ran our numerical method for 420 random communities for each integer value of $S$ between 2 and 100, and for all multiples of 10 between 100 and 1000 – and that is why we used two different colors in the plots – resulting in 79,380 solutions evaluated for each parameter set, from which no more than 100 errors were observed. In each panel, a different combination of the distributions from Fig. S1 was considered, with their specific values displayed within each panel. No clear tendency emerges from the plots.

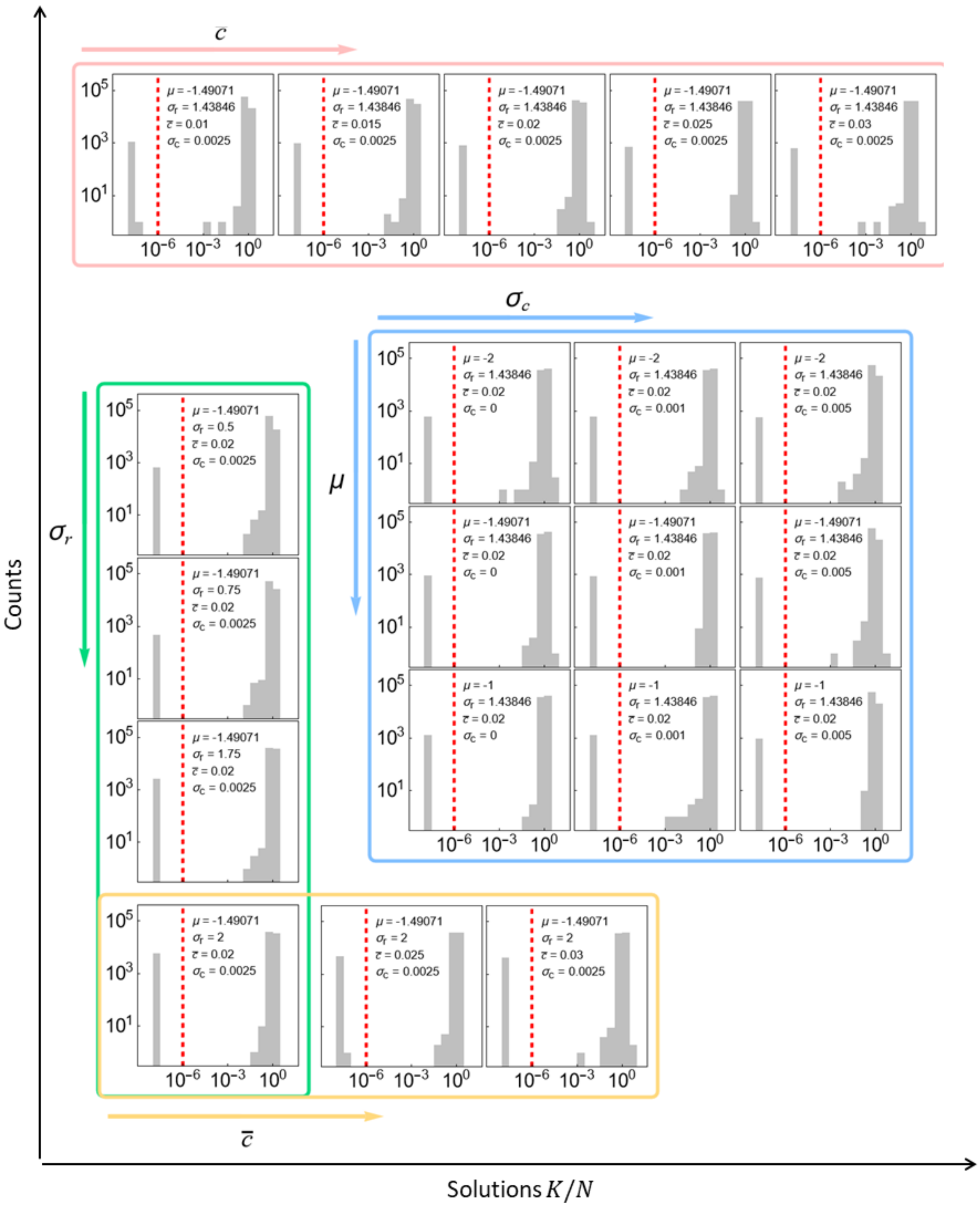


**Figure S3. There is a clear separation between the feasible and non-feasible solutions of the OFS.** In the panels, we show histograms of the solutions of Eq.(11) from the main text for $K/N$, where we can see a clear separation between values accumulating around $\sim 1$ (corresponding to feasible solutions) and values close to zero (corresponding to the non-feasible solutions, see Section S8). We notice that our chosen cutoff $K/N = 10^{-6}$ (dashed red line) is a good numerical threshold value to separate between the feasible and the non-feasible solutions.

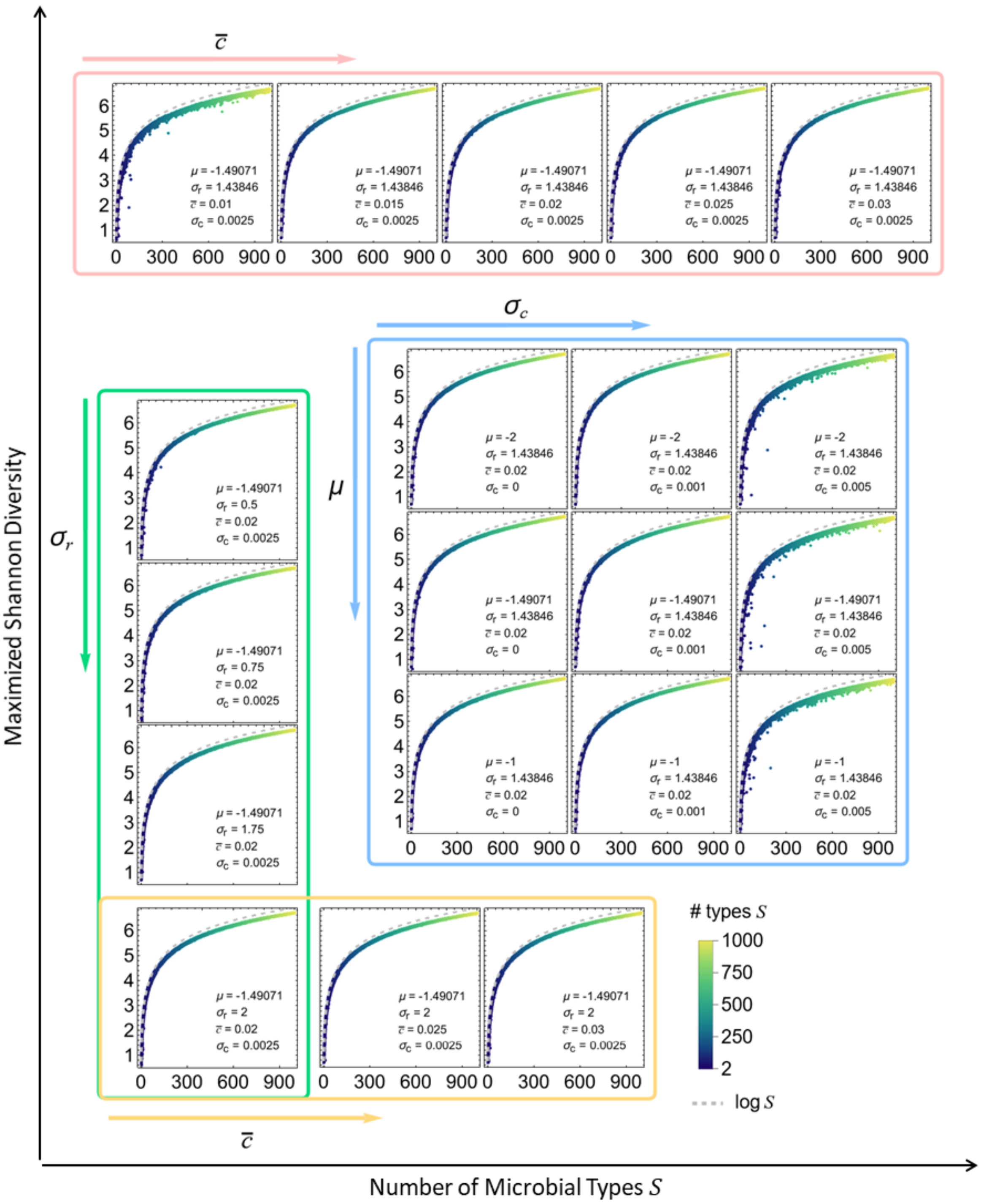


**Figure S4. The obtained maximized Shannon diversity preserves the same pattern throughout a large range of growth and clearance rate parameters.** In each panel, we plot the obtained maximal Shannon diversity whenever the solution of Eq.(11) from the main text is feasible, as a function of the number of microbial types $S$ (also identified by the color code). For a community with $S$ types, the maximal Shannon diversity is given by $\log S$, which is obtained when all types have the same abundance. Thus we plot this curve for comparison (the dashed gray line).

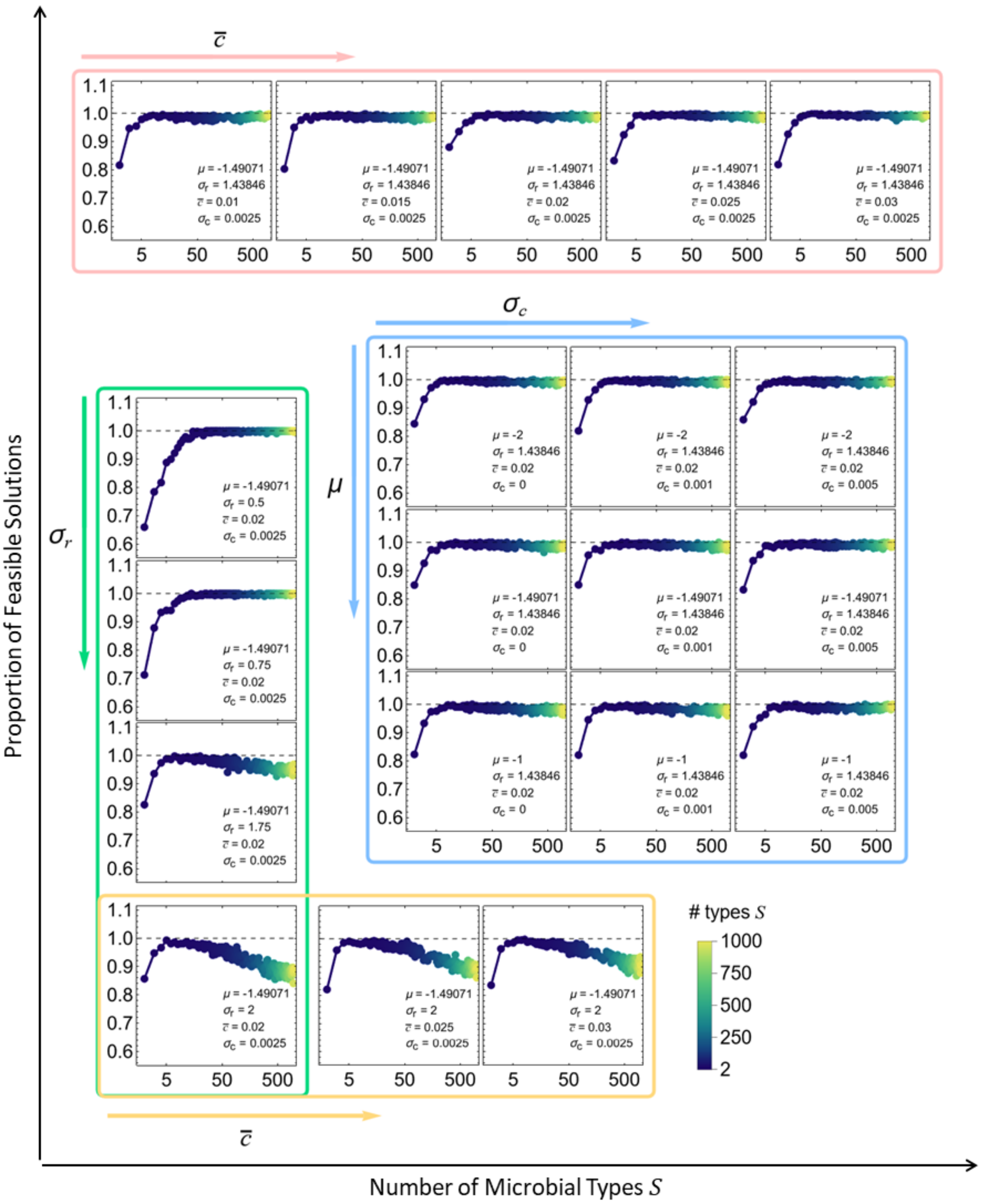


**Figure S5. The proportion of times an OFS exists grows fast to 1 for a large set of growth and clearance rate parameters.** In each panel, we plot the obtained proportion of feasible solutions as a function of the number of microbial types $S$ (also shown according to the color code). Apart from a specific region of the parameter sets ($r_i$ typically smaller than $c_i$, as discussed in the Results section of the main text), the existence of an OFS seems to be almost certain for communities with a large number of types.

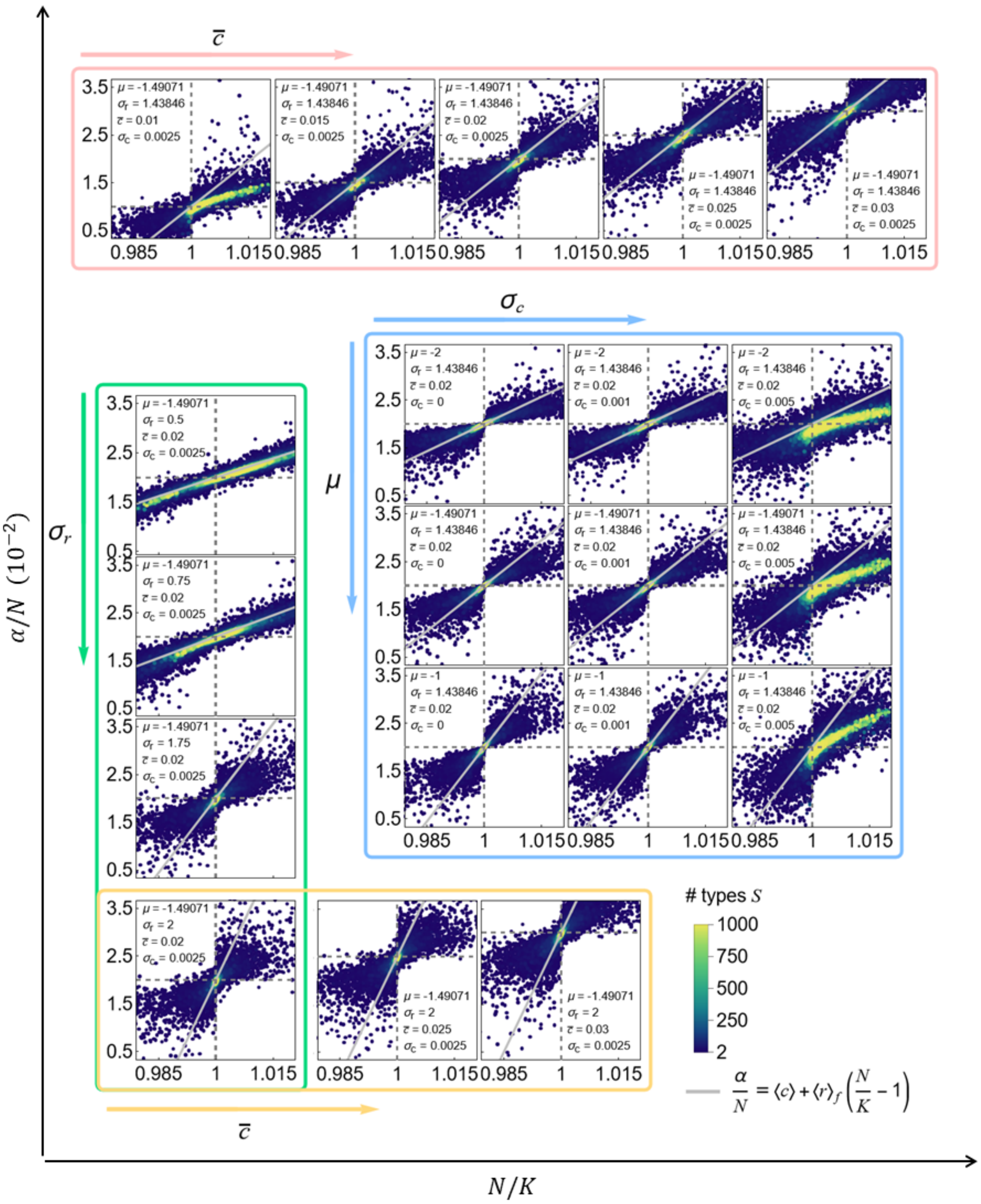

**Figure S6. The optimal feeding rate is fairly explained by the ansatz solution when the number of microbial types is large.** In each panel, all obtained feasible solution pairs $(N/K, \alpha/N)$ are plotted, with the color code standing for the number of types $S$. As $S$ increases, the solutions approach the ansatz (displayed as the gray continuous line), and, ultimately, the point $(N/K = 1, \alpha/N = \langle c \rangle)$. As discussed in the Results section of the main text, the convergence to the ansatz is slower for broader clearance rate distributions.

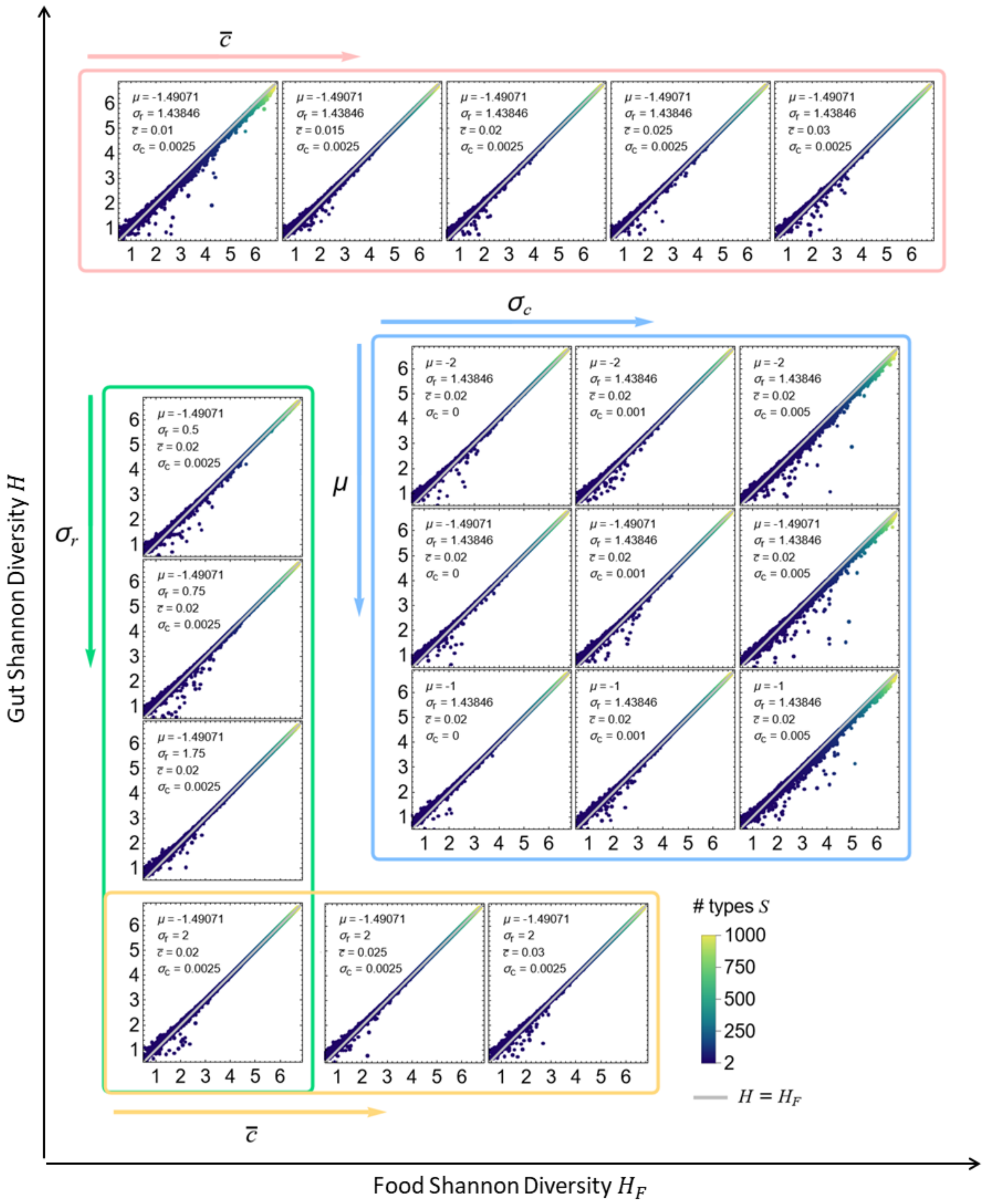


**Figure S7. The gut Shannon diversity converges to the food diversity when the number of microbial types is large.** In each panel, the maximized Shannon diversity $H$ in all feasible cases is plotted as a function of the corresponding Shannon diversity of the microbes in the food, $H_F$. We can observe that, for small numbers of microbial types $S$ (displayed as the color code), $H$ is generally higher than $H_F$. Indeed, for $S = 2$, the optimal parameters lead to $H = \log 2$. But as the number of types increases, $H$ converges to $H_F$ (with the curve $H = H_F$ plotted as the continuous gray line), a result observed for all simulated parameters, in line with the ansatz solution.

#### S2 Linear Stability Analysis of the abundance equations in the absence of feeding

We consider the system of equations

$$\frac{dn_i}{dt} = n_i r_i \left(1 - \frac{N}{K}\right) - c_i n_i = f_i(n_1, ..., n_S) \text{ for } i = 1, ..., S, \tag{S1}$$

In the general case, we assume all the $\frac{c_i}{r_i}$ to be different from one another. The non-trivial fixed points of the system are thus

$$n_i = \begin{cases} 0, \text{ for } i \neq i^* \\ N = K\left(1 - \frac{c_{i^*}}{r_{i^*}}\right) \end{cases}, \text{ for all } i^* \in \{1, ..., S\}. \tag{S2}$$

The Jacobian matrix of the system writes

$$\mathbb{J}_{ij} = \frac{\partial f_i}{\partial n_j} = \delta_{ij}\left(r_i - c_i - r_i \sum_j \frac{n_j}{K}\right) - \frac{n_i r_i}{K}, \tag{S3}$$

where $\delta_{ij}$ is the Kronecker's Delta. At a non-trivial fixed point $\mathbf{n}^* = \{0, \ldots, n_{i^*}, \ldots, 0\}$ the Jacobian becomes

$$\mathbb{J}(\mathbf{n}^*) = \begin{bmatrix} \left(\frac{r_1 c_{i^*}}{r_{i^*}} - c_1\right) & 0 & 0 & \cdots & 0 & \cdots & 0 \\ 0 & \left(\frac{r_2 c_{i^*}}{r_{i^*}} - c_2\right) & 0 & \cdots & 0 & \cdots & 0 \\ 0 & 0 & \left(\frac{r_3 c_{i^*}}{r_{i^*}} - c_3\right) & \cdots & 0 & \cdots & 0 \\ \vdots & \vdots & \vdots & \ddots & \vdots & & \vdots \\ -\frac{r_{i^*}N}{K} & -\frac{r_{i^*}N}{K} & -\frac{r_{i^*}N}{K} & \cdots & -\frac{r_{i^*}N}{K} & \cdots & -\frac{r_{i^*}N}{K} \\ \vdots & \vdots & \vdots & & \vdots & \ddots & \vdots \\ 0 & 0 & 0 & \cdots & 0 & \cdots & \left(\frac{r_S c_{i^*}}{r_{i^*}} - c_S\right) \end{bmatrix}, \tag{S4}$$

and the characteristic polynomial is then calculated as the determinant $|\mathbb{J}(\mathbf{n}^*) - \lambda \mathbb{I}_S|$, which, via cofactor expansion, is given by

$$p(\lambda) = -(r_{i^*} - c_{i^*} + \lambda) \prod_{i \neq i^*} \left[\left(\frac{r_i c_{i^*}}{r_{i^*}} - c_i\right) - \lambda\right] \tag{S5}$$

Hence, as long as there is a type $j \neq i^*$ such that $\frac{c_j}{r_j} < \frac{c_{i^*}}{r_{i^*}}$, there is going to be at least one positive eigenvalue $\lambda = r_j\left(\frac{c_{i^*}}{r_{i^*}} - \frac{c_j}{r_j}\right) > 0$. But if $i^*$ is chosen such that $c_{i^*}/r_{i^*}$ is the smallest over all types, and smaller than 1, such that $r_{i^*} > c_{i^*}$, then all the eigenvalues are negative. Therefore, the only stable fixed point in the system is the one that corresponds to all types being extinct but the one with the smallest ratio $c_{i^*}/r_{i^*}$. Therefore, the effective carrying capacity of the system is given by $N^* \equiv K\left(1 - \min_i \frac{c_{i^*}}{r_{i^*}}\right)$.

The typical time scale $T$ to reach the stable fixed point is set by $-1/\lambda_{\min}$, where $\lambda_{\min}$ is the least negative eigenvalue,

$$\lambda_{\min} = -\min\left[\{r_{i^*} - c_{i^*}\} \cup \left\{c_i - r_i \frac{c_{i^*}}{r_{i^*}}\right\}_{i \neq i^*}\right]. \tag{S6}$$

Hence, the effect of feeding on the community composition is bound to depend on how the feeding interval $\tau$ compares to $T$.

#### S3 The convexity of the total abundance: an important lemma

In this section, we present a useful lemma concerning the behaviur of the total abundance: In the long time limit $t \to \infty$, between two consecutive feeding events, the total abundance $N(t)$ is a convex decreasing function.

First, consider $\frac{dN}{dt}$,

$$\frac{dN}{dt} = \sum_i n_i \left(r_i - c_i - r_i \frac{N}{K}\right), \tag{S7}$$

thus the second derivative of $N(t)$ is given by

$$\begin{aligned} \frac{d^2N}{dt^2} &= \sum_i \frac{dn_i}{dt}\left(r_i - c_i - r_i\frac{N}{K}\right) - \sum_i \frac{n_i r_i}{k}\frac{dN}{dt} \\ &= \sum_i n_i \left(r_i - c_i - r_i\frac{N}{K}\right)^2 - \frac{dN}{dt}\sum_i \frac{n_i r_i}{K}. \end{aligned} \tag{S8}$$

Therefore, if $\frac{dN}{dt} < 0$, i.e., if the total abundance decreases, $N(t)$ is a convex function, i.e. $\frac{d^2N}{dt^2} > 0$.

Now, from Eq.(S7), if $N/K > 1 - c_i/r_i$ for every $i$, then $\frac{dN}{dt} < 0$, and therefore $N(t)$ decreases whenever $N/K > 1 - \min_i \frac{c_i}{r_i}$. But $N = K(1 - \min_i \frac{c_i}{r_i}) = N^*$ is the value of $N(t)$ at the stable fixed point in the absence of feeding. Hence, whenever the total abundance is larger than the effective carrying capacity $N^*$, $N(t)$ is a decreasing convex function.

Now, suppose that the system is close to its fixed point and some food is added at time $t^*$. Therefore, $N((t^*)^+) \approx N^* + n_f$ which is larger than $N^*$ and therefore right after this feeding event, $N(t)$ decreases in a convex way. If the following feeding event happens soon enough, i.e., $\tau < \tau^*$ for some $\tau^*$ to be determined, $N((t^* + \tau)^-)$ is still larger than $N^*$. As a consequence, $N(t)$ between two feeding events is a convex decreasing function for $\tau < \tau^*$. Taking $\tau^*$ as the fastest time scale of the dynamics is enough to guarantee $dN/dt$ did not change signs, i.e. $\tau^* = -1/\lambda_{\max}$ with

$$\lambda_{\max} = -\max\left[\{r_{i^*} - c_{i^*}\} \cup \left\{c_i - r_i \frac{c_{i^*}}{r_{i^*}}\right\}_{i \neq i^*}\right]. \tag{S9}$$

#### S4 The error of the trapezoidal approximation to the average total abundance

If $N(t)$ is a convex decreasing function (as in Figure S8a), we can write an upper bound to the error of $\langle N \rangle_\tau$ as

$$0 < E < n_f/2. \tag{S10}$$

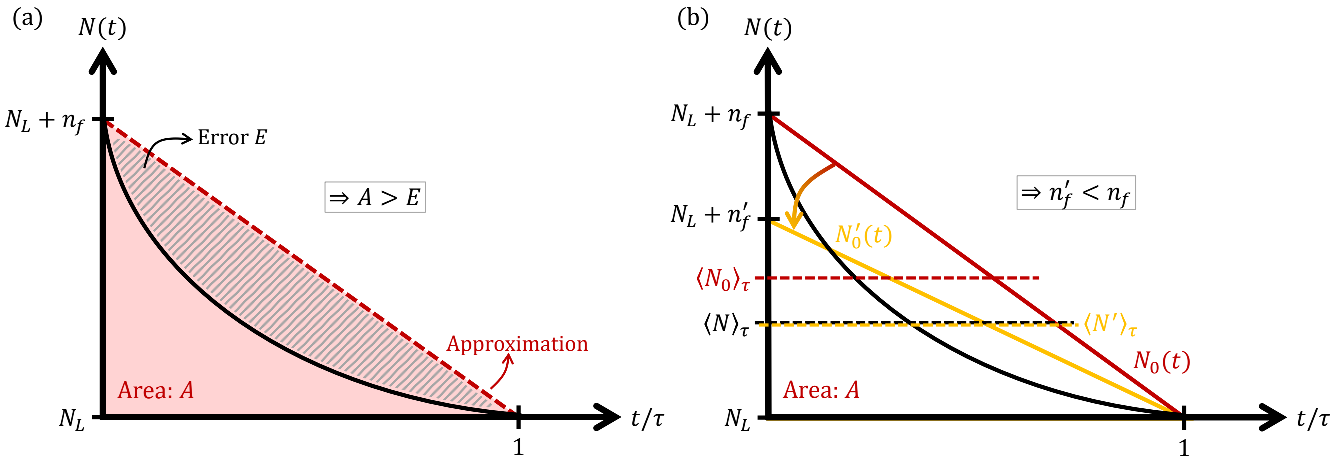


**Figure S8. The error of the trapezoidal approximation and the argument behind the existence condition for the OFS.** In (a), the graph of the total abundance $N$ as a function of time taken in units of $\tau$ (black continuous curve), is plotted with its trapezoidal approximation (red dashed curve). This approximation is used to calculate the integral of $N(t)$ divided by $\tau$, thus the error $E$ of this procedure is given by the hatched area. If $N(t)$ is a convex decreasing function between two feeding intervals (in the figure $t = 0$ and $t = \tau$), then $E$ is smaller than the area $A$ of the read triangle in the background. In (b), in addition to the curves $N(t)$ (in black) and its trapezoidal approximation $N_0(t)$ (in red), we plotted, in dashed lines, their average values, $\langle N\rangle_\tau$ and $\langle N_0\rangle_\tau$, respectively. Because of the decreasing convex shape of $N(t)$, $\langle N\rangle_\tau < \langle N_0\rangle_\tau$, thus leading to a different diversity value than the one realized without any approximation. Therefore, in order to obtain the same diversity, one needs to decrease the slope of $N_0(t)$, thus making the average $\langle N_0'\rangle_\tau$ of the new curve $N_0'(t)$ to coincide with $\langle N\rangle_\tau$, i.e., going from the red line to the yellow line in the figure. Notice that, as we define the OFS as a function $n_f = n_f(\tau)$, for each $\tau$, it corresponds to a smaller $n_f$ value, called $n_f'$ in the figure. And thus the linear OFS approximation (which emerges from the trapezoidal approximation in the continuous limit) results in an curve below the non-approximated OFS.

Now, if we write $\langle N\rangle_\tau = \langle N\rangle_\tau^{(0)} - E$, it results from Eq.(2) in the main text that

$$n_i(t) = n_i(t_0)\exp\left[\tau(r_i - c_i) - \frac{r_i\tau\langle N\rangle_\tau^{(0)}}{K} + \frac{r_i\tau E}{K}\right], \tag{S11}$$

hence we ask for $r_i\tau E/K \ll 1$ for every $i$. Therefore,

$$\max_i r_i \frac{\tau n_f}{2K} \ll 1 \tag{S12}$$

is a condition for the trapezoidal approximation to hold.

#### S5 Showing that a feasible linear OFS is a sufficient condition to the existence of a feasible OFS

Notice that the abundances $n_i(t)$ depend on the time-averaged total abundance $\langle N\rangle_\tau$, and so does the Shannon diversity. In Figure S8b, the black curve represents the total abundance $N(t)$ between two feeding events, and the red curve is its trapezoidal approximation $N_0(t)$. Considering $N(t)$ to be a decreasing convex function, then its average $N(t)$ is smaller than the average of $N_0(t)$. Hence, for a given $\tau$, a trapezoidal approximation that gives the same average $\langle N(t)\rangle_\tau$ considers a different (and unique) value $n'_f$, which is smaller than $n_f$. This fact explains why the linear approximation of the OFS lies below the true OFS in the heatmap of Fig. 2 from the main text.

The first consequence is that, if the linear approximation is a line with a positive slope, i.e., is feasible, then the OFS is never going to cross the $n_f = 0$ line for a positive value of $\tau$. Moreover, if the linear approximation is uniquely determined (as shown later in Section S8), since the $n'_f$ value is unique, the true OFS is indeed described by a function $n_f = n_f(\tau)$. Therefore, the existence of a feasible linear approximation of an OFS is a sufficient condition for the existence of a feasible OFS.

#### S6 Approximations of the time-averaged Shannon diversity

In this work, we consider the time average of the Shannon diversity in a specific form. Here we show how this form is a good approximation of other averaging choices, up to first order in small deviations around the absolute abundances. Let us start with the definition of the average Shannon diversity

$$\langle H\rangle_\tau = -\frac{1}{\tau}\int_{t_0}^{t_0+\tau}\sum_i \frac{n_i(t)}{N(t)}\log\frac{n_i(t)}{N(t)}dt = \left\langle \sum_i \frac{n_i(t)}{N(t)}\log\frac{n_i(t)}{N(t)}\right\rangle_\tau \tag{S13}$$

for $t_0 \to \infty$.

Now, let

$$n_i(t) = \langle n_i\rangle_\tau + \delta n_i(t) \text{ and } N(t) = \langle N\rangle_\tau + \delta N(t), \tag{S14}$$

which defines the deviation functions $\delta n_i(t)$ and $\delta N(t)$, satisfying $\langle \delta N(t)\rangle_\tau = \langle \delta n_i(t)\rangle_\tau = 0$. We expect those deviations to be small when $n_f\tau/K \ll 1$. With this definition, first we calculate the fraction $n_i/N$ as

$$\begin{aligned}
\frac{n_i}{N} &= \frac{\langle n_i\rangle_\tau + \delta n_i}{\langle N\rangle_\tau + \delta N}\\
&= \frac{\langle n_i\rangle_\tau}{\langle N\rangle_\tau}\left(1+\frac{\delta n_i}{\langle n_i\rangle_\tau}\right)\left(1+\frac{\delta N}{\langle N\rangle_\tau}\right)^{-1}\\
&= \frac{\langle n_i\rangle_\tau}{\langle N\rangle_\tau}\left(1+\frac{\delta n_i}{\langle n_i\rangle_\tau}\right)\left(1-\frac{\delta N}{\langle N\rangle_\tau}+\frac{(\delta N)^2}{\langle N\rangle_\tau^2}+O(\delta^3)\right)\\
&= \frac{\langle n_i\rangle_\tau}{\langle N\rangle_\tau}\left(1-\frac{\delta N}{\langle N\rangle_\tau}+\frac{\delta n_i}{\langle n_i\rangle_\tau}+\frac{(\delta N)^2}{\langle N\rangle_\tau^2}-\frac{\delta N\delta n_i}{\langle N\rangle_\tau\langle n_i\rangle_\tau}+O(\delta^3)\right),
\end{aligned} \tag{S15}$$

where we used that $(1+x)^{-1} = 1-x+x^2+O(x^3)$, and wrote $\delta$ as a generic term for any deviation. Now, for $\log(n_i/N)$,

$$\begin{aligned}
\log\frac{n_i}{N} &= \log\frac{\langle n_i\rangle_\tau}{\langle N\rangle_\tau}+\log\left(1-\frac{\delta N}{\langle N\rangle_\tau}+\frac{\delta n_i}{\langle n_i\rangle_\tau}+\frac{(\delta N)^2}{\langle N\rangle_\tau^2}-\frac{\delta N\delta n_i}{\langle N\rangle_\tau\langle n_i\rangle_\tau}+O(\delta^3)\right)\\
&= \log\frac{\langle n_i\rangle_\tau}{\langle N\rangle_\tau}-\frac{\delta N}{\langle N\rangle_\tau}+\frac{\delta n_i}{\langle n_i\rangle_\tau}+\frac{(\delta N)^2}{\langle N\rangle_\tau^2}-\frac{\delta N\delta n_i}{\langle N\rangle_\tau\langle n_i\rangle_\tau}-\frac{1}{2}\left(\frac{(\delta N)^2}{\langle N\rangle_\tau^2}+\frac{(\delta n_i)^2}{\langle n_i\rangle_\tau^2}-2\frac{\delta N\delta n_i}{\langle N\rangle_\tau\langle n_i\rangle_\tau}\right)+O(\delta^3)\\
&= \log\frac{\langle n_i\rangle_\tau}{\langle N\rangle_\tau}-\frac{\delta N}{\langle N\rangle_\tau}+\frac{\delta n_i}{\langle n_i\rangle_\tau}+\frac{(\delta N)^2}{\langle N\rangle_\tau^2}-\frac{\delta N\delta n_i}{\langle N\rangle_\tau\langle n_i\rangle_\tau}-\frac{1}{2}\left(\frac{\delta N}{\langle N\rangle_\tau}-\frac{\delta n_i}{\langle n_i\rangle_\tau}\right)^2+O(\delta^3),
\end{aligned} \tag{S16}$$

where we used that $\log(1+x) = x - x^2/2 + O(x^3)$. Now, combining both results,

$$\langle H\rangle_\tau = -\sum_i \frac{\langle n_i\rangle_\tau}{\langle N\rangle_\tau}\left\langle \left(1 - \frac{\delta N}{\langle N\rangle_\tau} + \frac{\delta n_i}{\langle n_i\rangle_\tau} + \frac{(\delta N)^2}{\langle N\rangle_\tau^2} - \frac{\delta N \delta n_i}{\langle N\rangle_\tau \langle n_i\rangle_\tau} + O(\delta^3)\right)\right.$$
$$\left. \times \left(\log\frac{\langle n_i\rangle_\tau}{\langle N\rangle_\tau} - \frac{\delta N}{\langle N\rangle_\tau} + \frac{\delta n_i}{\langle n_i\rangle_\tau} + \frac{(\delta N)^2}{\langle N\rangle_\tau^2} - \frac{\delta N \delta n_i}{\langle N\rangle_\tau \langle n_i\rangle_\tau} - \frac{1}{2}\left(\frac{\delta N}{\langle N\rangle_\tau} - \frac{\delta n_i}{\langle n_i\rangle_\tau}\right)^2 + O(\delta^3)\right)\right\rangle$$
$$= -\sum_i \frac{\langle n_i\rangle_\tau}{\langle N\rangle_\tau}\left\langle \log\frac{\langle n_i\rangle_\tau}{\langle N\rangle_\tau} + \frac{(\delta N)^2}{\langle N\rangle_\tau^2} - \frac{\delta N \delta n_i}{\langle N\rangle_\tau \langle n_i\rangle_\tau} + \frac{1}{2}\left(\frac{\delta N}{\langle N\rangle_\tau} - \frac{\delta n_i}{\langle n_i\rangle_\tau}\right)^2 \right.$$
$$\left. + \left(\frac{(\delta N)^2}{\langle N\rangle_\tau^2} - \frac{\delta N \delta n_i}{\langle N\rangle_\tau \langle n_i\rangle_\tau}\right)\log\frac{\langle n_i\rangle_\tau}{\langle N\rangle_\tau}\right\rangle + O(\delta^3). \tag{S17}$$

Using that $\sum_i \delta n_i = \delta N$ and that $\sum_i \langle n_i\rangle_\tau = \langle N\rangle_\tau$, we get

$$\sum_i \frac{\langle n_i\rangle_\tau}{\langle N\rangle_\tau}\left\langle \frac{(\delta N)^2}{\langle N\rangle_\tau^2} - \frac{\delta N \delta n_i}{\langle N\rangle_\tau \langle n_i\rangle_\tau}\right\rangle = 0. \tag{S18}$$

Hence,

$$\langle H\rangle_\tau = \langle H\rangle_\tau^{(1)} - \frac{1}{2}\sum_i \frac{\langle n_i\rangle_\tau}{\langle N\rangle_\tau}\left\langle \left(\frac{\delta N}{\langle N\rangle_\tau} - \frac{\delta n_i}{\langle n_i\rangle_\tau}\right)^2\right\rangle_\tau - \sum_i \frac{\langle n_i\rangle_\tau}{\langle N\rangle_\tau}\log\frac{\langle n_i\rangle_\tau}{\langle N\rangle_\tau}\left(\frac{\langle(\delta N)^2\rangle}{\langle N\rangle_\tau^2} - \frac{\langle \delta N \delta n_i\rangle}{\langle N\rangle_\tau \langle n_i\rangle_\tau}\right) + O(\delta^3), \tag{S19}$$

where

$$\langle H\rangle_\tau^{(1)} \equiv -\sum_i \frac{\langle n_i\rangle_\tau}{\langle N\rangle_\tau}\log\frac{\langle n_i\rangle_\tau}{\langle N\rangle_\tau} \tag{S20}$$

is the approximation we used in this work. Eq.(S19) shows that $\langle H\rangle_\tau^{(1)} \approx \langle H\rangle_\tau$ is a good approximation up to the first order of small abundance deviations from the average, as it also calculates the second order corrections.

A second natural approximation of the average diversity is given by

$$\langle H\rangle_\tau^{(2)} \equiv -\sum_i \left\langle \frac{n_i}{N}\right\rangle_\tau \log\left\langle \frac{n_i}{N}\right\rangle_\tau, \tag{S21}$$

and we show now that this is also a good approximation up to first order deviations of the absolute abundances. Taking the time average of Eq.(S15), we find

$$\left\langle \frac{n_i}{N}\right\rangle_\tau = \frac{\langle n_i\rangle_\tau}{\langle N\rangle_\tau}\left(1 + \frac{\langle(\delta N)^2\rangle_\tau}{\langle N\rangle_\tau^2} - \frac{\langle \delta N \delta n_i\rangle_\tau}{\langle N\rangle_\tau \langle n_i\rangle_\tau} + O(\delta^3)\right), \tag{S22}$$

and then, for $\log\left\langle \frac{n_i}{N}\right\rangle_\tau$, we find

$$\log\left\langle \frac{n_i}{N}\right\rangle_\tau = \log\frac{\langle n_i\rangle_\tau}{\langle N\rangle_\tau} + \frac{\langle(\delta N)^2\rangle_\tau}{\langle N\rangle_\tau^2} - \frac{\langle \delta N \delta n_i\rangle_\tau}{\langle N\rangle_\tau \langle n_i\rangle_\tau} + O(\delta^3). \tag{S23}$$

Combining both equations,

$$\langle H\rangle_\tau^{(2)} = -\sum_i \frac{\langle n_i\rangle_\tau}{\langle N\rangle_\tau}\left(\log\frac{\langle n_i\rangle_\tau}{\langle N\rangle_\tau} + \frac{\langle(\delta N)^2\rangle_\tau}{\langle N\rangle_\tau^2} - \frac{\langle \delta N \delta n_i\rangle_\tau}{\langle N\rangle_\tau\langle n_i\rangle_\tau} + \log\frac{\langle n_i\rangle_\tau}{\langle N\rangle_\tau}\left(\frac{\langle(\delta N)^2\rangle_\tau}{\langle N\rangle_\tau^2} - \frac{\langle \delta N \delta n_i\rangle_\tau}{\langle N\rangle_\tau\langle n_i\rangle_\tau}\right)\right) + O(\delta^3)$$

$$= \langle H\rangle_\tau^{(1)} - \sum_i \frac{\langle n_i\rangle_\tau}{\langle N\rangle_\tau}\log\frac{\langle n_i\rangle_\tau}{\langle N\rangle_\tau}\left(\frac{\langle(\delta N)^2\rangle_\tau}{\langle N\rangle_\tau^2} - \frac{\langle \delta N \delta n_i\rangle_\tau}{\langle N\rangle_\tau\langle n_i\rangle_\tau}\right) + O(\delta^3). \quad \text{(S24)}$$

Subtracting $\langle H\rangle_\tau$ from this last equation, we finally get

$$\langle H\rangle_\tau = \langle H\rangle_\tau^{(2)} - \frac{1}{2}\sum_i \frac{\langle n_i\rangle_\tau}{\langle N\rangle_\tau}\left\langle\left(\frac{\delta N}{\langle N\rangle_\tau} - \frac{\delta n_i}{\langle n_i\rangle_\tau}\right)^2\right\rangle_\tau + O(\delta^3), \quad \text{(S25)}$$

as we wanted to show.

To summarize these results, let us define

$$\begin{cases} \Delta h_1 \equiv -\frac{1}{2}\sum_i \frac{\langle n_i\rangle_\tau}{\langle N\rangle_\tau}\left\langle\left(\frac{\delta N}{\langle N\rangle_\tau} - \frac{\delta n_i}{\langle n_i\rangle_\tau}\right)^2\right\rangle_\tau \\ \Delta h_2 \equiv -\sum_i \frac{\langle n_i\rangle_\tau}{\langle N\rangle_\tau}\log\frac{\langle n_i\rangle_\tau}{\langle N\rangle_\tau}\left(\frac{\langle(\delta N)^2\rangle_\tau}{\langle N\rangle_\tau^2} - \frac{\langle \delta N \delta n_i\rangle_\tau}{\langle N\rangle_\tau\langle n_i\rangle_\tau}\right) \end{cases}, \quad \text{(S26)}$$

and then

$$\begin{cases} \langle H\rangle_\tau = \langle H\rangle_\tau^{(1)} + \Delta h_1 + \Delta h_2 + O(\delta^3) \\ \langle H\rangle_\tau = \langle H\rangle_\tau^{(2)} + \Delta h_1 + O(\delta^3) \end{cases}. \quad \text{(S27)}$$

It is easy to see that $\Delta h_1 \leq 0$ and therefore

$$\langle H\rangle_\tau \leq \langle H\rangle_\tau^{(2)}, \quad \text{(S28)}$$

which agrees with the Jensen's inequality for concave functions like the Shannon entropy, $\langle H(\{\rho_i\})\rangle \leq H(\{\langle\rho_i\rangle\})$.

Figure S9 shows these different results for different feeding parameters in an $S = 2$ case.

#### S7 Maximizing the Shannon diversity in the continuous feeding limit for $S > 2$

The problem can be stated as: Given a diversity index $H = H(\alpha, N)$, with $n_i = n_i(\alpha, N)$, find its maximum subjected to the constraint $n_1 + \ldots + n_S = N$. $H$ is the Shannon diversity and we use Lagrange multipliers to solve this optimization problem,

$$\begin{cases} n_i = \dfrac{\alpha f_i}{c_i - r_i + r_i N/K}, \\ H = -\sum_i (n_i/N)\log(n_i/N) = H(\alpha, N), \\ \sum_{i=1}^S n_i = N, \end{cases} \quad \text{(S29)}$$

with the Lagrangian

$$\mathscr{L}(\alpha, N, \lambda) = H(\alpha, N) - \lambda\left(\sum_{i=1}^S n_i - N\right). \quad \text{(S30)}$$

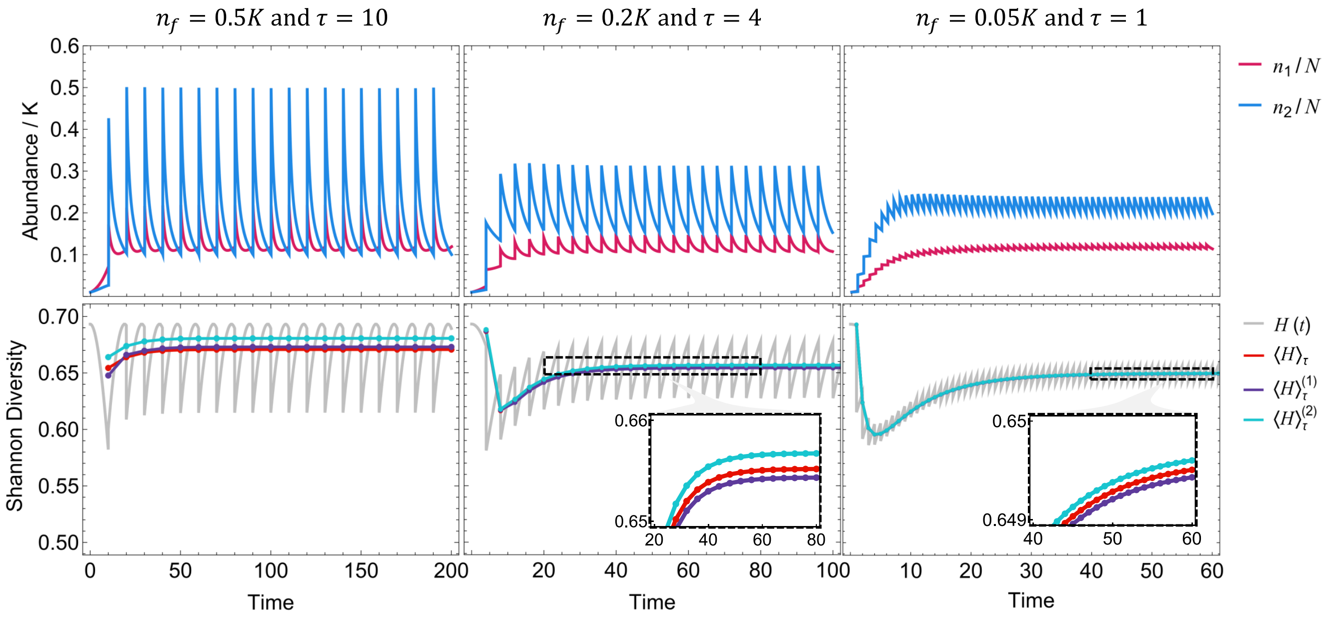


**Figure S9. The different approximations of the time-averaged Shannon Diversity converge when $\tau n_f/K \to 0$.** In each column we numerically solve the model for a 2-types case, considering decreasing values of $n_f\tau/K$. In the first row, the abundances are plotted as a function of time, whereas in the second row, the corresponding instantaneous Shannon diversity $H(t)$ is shown in gray. The average of the previous time interval $\tau$, $\langle H \rangle_\tau$ is marked in red, whereas the two different approximations discussed in the text, $\langle H \rangle_\tau^{(1)}$ (given by Eq.(S20)) and $\langle H \rangle_\tau^{(2)}$ (given by Eq.(S21)), are plotted in purple and cyan, respectively.

We must solve $\nabla_{\alpha.N,\lambda}\mathscr{L} = 0$,

$$\frac{\partial \mathscr{L}}{\partial \alpha} = \sum_i \frac{\partial H}{\partial n_i}\frac{\partial n_i}{\partial \alpha} - \lambda \sum_i \frac{\partial n_i}{\partial \alpha} = 0, \tag{S31}$$

$$\frac{\partial \mathscr{L}}{\partial N} = -\sum_i \left(\log(n_i/N)+1\right)\frac{\partial}{\partial N}\left(\frac{n_i}{N}\right) - \lambda\left(\sum_i \frac{\partial n_i}{\partial N} - 1\right) = 0, \tag{S32}$$

$$\frac{\partial \mathscr{L}}{\partial \lambda} = \sum_i n_i - N = 0. \tag{S33}$$

Calculating the derivatives:

$$\frac{\partial n_i}{\partial \alpha} = n_i/\alpha, \tag{S34}$$

$$\frac{\partial n_i}{\partial N} = -\frac{\alpha f_i r_i/K}{(c_i - r_i + r_i N/K)^2} = -\frac{n_i^2 r_i}{K\alpha f_i}, \tag{S35}$$

$$\frac{\partial}{\partial N}\left(\frac{n_i}{N}\right) = -\frac{n_i^2 r_i}{NK\alpha f_i} - \frac{n_i}{N^2} = -\frac{1}{N}\left(\frac{n_i^2 r_i}{K\alpha f_i} + \frac{n_i}{N}\right), \tag{S36}$$

$$\frac{\partial H}{\partial n_i} = -\frac{1}{N}\left(\log(n_i/N)+1\right). \tag{S37}$$

And now, combining equations (S31) - (S37), we find

$$\sum_i n_i \frac{\partial H}{\partial n_i} = \lambda N \Rightarrow H - 1 = \lambda N, \tag{S38}$$

and for the diversity,

$$H = -\frac{\sum_i \frac{n_i^2 r_i}{f_i} \log(n_i/N)}{\sum_i \frac{n_i^2 r_i}{f_i}}. \tag{S39}$$

We define now,

$$n_i/N \equiv \rho_i \equiv (\alpha/N)\widetilde{\rho}_i, \tag{S40}$$

in which $\widetilde{\rho}_i = f_i/(c_i - r_i + r_i N/K)$ does not depend explicitly on $\alpha$. From the definition of $H$, we can write

$$\begin{aligned} H &= -\sum_i \rho_i \log \rho_i \\ &= -(\alpha/N)\sum_i \widetilde{\rho}_i \log((\alpha/N)\widetilde{\rho}_i) \\ &= -(\alpha/N)\sum_i \widetilde{\rho}_i (\log(\alpha/N) + \log \widetilde{\rho}_i) \\ &= -\log(\alpha/N) - (\alpha/N)\sum_i \widetilde{\rho}_i \log \widetilde{\rho}_i; \end{aligned} \tag{S41}$$

and from the result in equation (S39),

$$\begin{aligned} H &= -\frac{\sum_i (\widetilde{\rho}_i^2 r_i/f_i)\log((\alpha/N)\widetilde{\rho}_i)}{\sum_i (\widetilde{\rho}_i^2 r_i/f_i)} \\ &= -\log(\alpha/N) - \frac{\sum_i (\widetilde{\rho}_i^2 r_i/f_i)\log \widetilde{\rho}_i}{\sum_i (\widetilde{\rho}_i^2 r_i/f_i)}. \end{aligned} \tag{S42}$$

Finally, combining both equations,

$$(\alpha/N) = \frac{\sum_i (\widetilde{\rho}_i^2 r_i/f_i)\log \widetilde{\rho}_i}{\sum_j (\widetilde{\rho}_j \log \widetilde{\rho}_j)\sum_i (\widetilde{\rho}_i^2 r_i/f_i)}, \tag{S43}$$

which, together with the constraint $(\alpha/N)\sum_i \widetilde{\rho}_i = 1$ solves the system for $\alpha/N$ and $N/K$ – which will be made clearer in the next section.

#### S8 The numerical method to solve the linear OFS approximation equations

A numerical solution for $\alpha$ can be obtained by combining Eq.(S43) with the constraint $(\alpha/N)\sum_i \widetilde{\rho}_i = 1$. However, a unique solution to this equation is not guaranteed, and in general, as the number of types increases, a numerical method can show problems to find all the existing solutions. In this section, we propose a fast numerical algorithm to find the (global) optimal solution of the system.

Multiplying Eq.(S43) by $\widetilde{\rho}_j$ and summing it over $j$, we obtain

$$1 = \frac{\left(\sum_j \widetilde{\rho}_j\right) \sum_i (\widetilde{\rho}_i^2 r_i / f_i) \log \widetilde{\rho}_i}{\sum_j (\widetilde{\rho}_j \log \widetilde{\rho}_j) \sum_i (\widetilde{\rho}_i^2 r_i / f_i)}. \tag{S44}$$

Multiplying both sides by the denominator of the right-hand side and rearranging all the terms to be on the same side, we find an equation for $N$ that does not depend on $\alpha$,

$$G(N) = \sum_{i,j} \frac{\widetilde{\rho}_i \widetilde{\rho}_j^2 r_j}{f_j} \log \frac{\widetilde{\rho}_i}{\widetilde{\rho}_j} = 0, \tag{S45}$$

whose roots can be searched for numerically. We are interested in communities where all types survive: indeed, without feeding, most microbial types will go extinct, but migration necessarily increases the average abundances of all types to positive values. Hence, $\rho_i$ should be positive for every $i$. Because of that,

$$N > K\left(1 - \min_i \frac{c_i}{r_i}\right) = N^*. \tag{S46}$$

Now, if $N^* > 0$,

$$\frac{\partial H}{\partial N} = -\frac{\alpha}{N^2} \sum_i f_i \frac{(c_i - r_i + 2r_i N/K)}{(c_i - r_i + r_i N/K)^2} |\log \rho_i| < 0 \tag{S47}$$

for $N > N^*$, i.e., the diversity decreases as $N$ increases. Therefore, the solution of equation (S45) that maximizes diversity is the smallest root of $G(N)$ that is bigger than $N^*$. Since $G(\{\widetilde{\rho}_i\})$ has a singularity at $N = N^*$, a numerical method to find the root of $G(N)$ (e.g. Newton's) that uses $(1 + |\varepsilon|)N^*$ as initial value (where $|\varepsilon| \ll 1$), can be effective to find the optimal solution.

Note that, as $N \to \infty$, $G(N) \to 0$, and thus in the absence of feasible solutions, this method can result in numerically blowing-up values of $N$. When comparing all the solutions we obtain, there is a clear separation between feasible solutions and such unrealistically large values, as shown in Figure S3. In practice, we used a numerical solver to find the roots of $G(N)$ in terms of $K/N$, and hence "blowing up" solutions are the ones where $K/N$ solutions are close to zero.

#### S9 An ansatz solution to the linear OFS approximation equations

Suppose that the optimal solution is close to the carrying capacity

$$N/K = 1 + \delta. \tag{S48}$$

We want to show that the diversity in the gut is given by

$$H = H_f - \left|\frac{d^2H}{d\delta^2}\right|_{\delta=0} \delta^2 + O(\delta^3) \tag{S49}$$

where $H_f = -\sum_i f_i \log f_i$ is the diversity in the food, showing that $H_f$ has a local maximum at $H = H_f$, which is achieved when $N/K = 1$.

We consider the case in which $c_i = c$ for every type $i$, and Taylor expand the diversity $H$ around $\delta = 0$,

$$H = H|_{\delta=0} + \left.\frac{dH}{d\delta}\right|_{\delta=0} \delta + \left.\frac{d^2H}{d\delta^2}\right|_{\delta=0} \delta^2 + O(\delta^3). \tag{S50}$$

First, the zeroth-order term:

$$\widetilde{\rho}_i = \frac{f_i}{c + r_i\delta} \Rightarrow \widetilde{\rho}_i|_{\delta=0} = f_i/c; \tag{S51}$$

$$(\alpha/N) = \frac{1}{\sum_i \widetilde{\rho}_i} \Rightarrow (\alpha/N)|_{\delta=0} = c; \tag{S52}$$

From Eq.(S41),

$$\begin{aligned} H = -\log(\alpha/N) - (\alpha/N)\sum_i \widetilde{\rho}_i \log\widetilde{\rho}_i \Rightarrow H|_{\delta=0} &= -\log c - c\sum_i \frac{f_i}{c}\log\frac{f_i}{c} \\ &= -\sum_i f_i \log f_i \\ &= H_f. \end{aligned} \tag{S53}$$

To finish the proof, we need to show that $dH/d\delta = 0$ and that $d^2H/d\delta^2 < 0$ when $\delta = 0$. For the first derivative term:

$$\frac{d\widetilde{\rho}_i}{d\delta} = \frac{-f_i r_i}{(c + r_i\delta)^2} \Rightarrow \left.\frac{d\widetilde{\rho}_i}{d\delta}\right|_{\delta=0} = \frac{-f_i r_i}{c^2}; \tag{S54}$$

$$\frac{d(\alpha/N)}{d\delta} = -\left(\sum_i \widetilde{\rho}_i\right)^{-2} \sum_i \frac{d\widetilde{\rho}_i}{d\delta} \Rightarrow \left.\frac{d(\alpha/N)}{d\delta}\right|_{\delta=0} = c^2 \sum_i \frac{f_i r_i}{c^2} = \langle r \rangle_f; \tag{S55}$$

$$\begin{aligned} \frac{dH}{d\delta} &= \frac{-1}{(\alpha/N)}\frac{d(\alpha/N)}{d\delta} - \frac{d(\alpha/N)}{d\delta}\sum_i \widetilde{\rho}_i \log\widetilde{\rho}_i - (\alpha/N)\sum_i \frac{d\widetilde{\rho}_i}{d\delta}(\log\widetilde{\rho}_i + 1) \\ &= -\frac{d(\alpha/N)}{d\delta}\sum_i \widetilde{\rho}_i(1 + \log\widetilde{\rho}_i) - (\alpha/N)\sum_i \frac{d\widetilde{\rho}_i}{d\delta}(\log\widetilde{\rho}_i + 1) \\ &= -\sum_i (\log\widetilde{\rho}_i + 1)\left(\widetilde{\rho}_i \frac{d(\alpha/N)}{d\delta} + (\alpha/N)\frac{d\widetilde{\rho}_i}{d\delta}\right) \\ &= -\sum_i \log\widetilde{\rho}_i \left(\widetilde{\rho}_i \frac{d(\alpha/N)}{d\delta} + (\alpha/N)\frac{d\widetilde{\rho}_i}{d\delta}\right) \end{aligned}$$

$$\Rightarrow \left.\frac{dH}{d\delta}\right|_{\delta=0} = -\sum_i \log\frac{f_i}{c}\left(\frac{f_i}{c}\langle r\rangle_f - c\frac{f_i r_i}{c^2}\right) = -\frac{1}{c}\left(\langle r\rangle_f\langle \log f\rangle_f + \langle r\log f\rangle_f\right). \tag{S56}$$

In the limit of an infinite number of types, $S \to \infty$, because $f_i$ and $r_i$ are uncorrelatedly drawn,

$$\langle r\rangle_f\langle \log f\rangle_f = \langle r\log f\rangle_f \tag{S57}$$

and then

$$\left.\frac{dH}{d\delta}\right|_{\delta=0} = 0. \tag{S58}$$

Now, for the second derivative,

$$\frac{d^2\widetilde{\rho}_i}{d\delta^2} = \frac{2f_i r_i^2}{(c+r_i\delta)^3} \Rightarrow \left.\frac{d^2\widetilde{\rho}_i}{d\delta^2}\right|_{\delta=0} = \frac{2f_i r_i^2}{c^3}; \tag{S59}$$

$$\frac{d^2(\alpha/N)}{d\delta^2} = 2\left(\sum_i \widetilde{\rho}_i\right)^{-3}\left(\sum_i \frac{d\widetilde{\rho}_i}{d\delta}\right)^2 - \left(\sum_i \widetilde{\rho}_i\right)^{-2}\sum_i \frac{d^2\widetilde{\rho}_i}{d\delta^2};$$

$$\Rightarrow \left.\frac{d^2(\alpha/N)}{d\delta^2}\right|_{\delta=0} = 2c^3\left(\sum_i \frac{f_i r_i}{c^2}\right)^2 - c^2\sum_i \frac{2f_i r_i^2}{c^3} = \frac{2}{c}\left(\langle r\rangle_f^2 - \langle r^2\rangle_f\right) = -\frac{2}{c}\mathrm{Var}_f(r). \tag{S60}$$

$$\frac{d^2H}{d\delta^2} = -\sum_i \frac{1}{\widetilde{\rho}_i}\frac{d\widetilde{\rho}_i}{d\delta}\left(\widetilde{\rho}_i\frac{d(\alpha/N)}{d\delta} + (\alpha/N)\frac{d\widetilde{\rho}_i}{d\delta}\right) - \sum_i \log\widetilde{\rho}_i\left(2\frac{d\widetilde{\rho}_i}{d\delta}\frac{d(\alpha/N)}{d\delta} + (\alpha/N)\frac{d^2\widetilde{\rho}_i}{d\delta^2} + \widetilde{\rho}_i\frac{d^2(\alpha/N)}{d\delta^2}\right) \tag{S61}$$

$$\begin{aligned}
\Rightarrow \left.\frac{d^2H}{d\delta^2}\right|_{\delta=0} &= -\sum_i \frac{c}{f_i}\frac{-f_i r_i}{c^2}\left(\frac{f_i}{c}\langle r\rangle_f + c\frac{-f_i r_i}{c^2}\right) - \sum_i \log\frac{f_i}{c}\left(2\frac{-f_i r_i}{c^2}\langle r\rangle_f + c\frac{2f_i r_i^2}{c^3} + \frac{f_i}{c}\frac{-2}{c}\mathrm{Var}_f(r)\right)\\
&= -\frac{1}{c^2}(\langle r^2\rangle_f - \langle r\rangle_f^2) + \frac{2}{c^2}\left(\langle r\log f\rangle_f\langle r\rangle_f - \langle r^2\log f\rangle_f + \langle \log f\rangle_f\mathrm{Var}_f(r)\right)\\
&= -\frac{1}{c^2}\mathrm{Var}_f(r) + \frac{2}{c^2}\langle \log f\rangle_f\left(\mathrm{Var}_f(r) - \frac{\langle r^2\log f\rangle_f - \langle r\log f\rangle_f\langle r\rangle_f}{\langle \log f\rangle_f}\right).
\end{aligned} \tag{S62}$$

As before, the term in parenthesis vanishes as $S \to \infty$, since in this limit any sampling correlation between $f_i$ and $r_i$ disappears and

$$\langle r\log f\rangle_f = \langle r\rangle_f\langle \log f\rangle_f \text{ and } \langle r^2\log f\rangle_f = \langle r^2\rangle_f\langle \log f\rangle_f. \tag{S63}$$

Therefore,

$$\left.\frac{d^2H}{d\delta^2}\right|_{\delta=0} = -\frac{1}{c^2}\mathrm{Var}_f(r) < 0. \tag{S64}$$

#### S10 The ansatz when the clearance rates are broadly distributed

Let us analyze now the maximal diversity $H$ when $c_i \equiv \langle c \rangle + \Delta c_i$. Now, $\rho_i|_{\delta=0} = (\alpha/K) f_i/c_i$ and then, from the constraint equation, we find

$$\begin{aligned}
\frac{K}{\alpha} &= \sum_i \frac{f_i}{c_i} \\
&= \sum_i \frac{f_i}{\langle c \rangle}\left(1 + \frac{\Delta c_i}{\langle c \rangle}\right)^{-1} \\
&= \frac{1}{\langle c \rangle}\left(1 - \sum_i f_i \frac{\Delta c_i}{\langle c \rangle}\right) + O((\Delta c_i/\langle c \rangle)^2). \qquad \text{(S65)}
\end{aligned}$$

And for the diversity $H = -\sum_i \rho_i \log \rho_i$,

$$\begin{aligned}
H|_{\delta=0} &= -\frac{\alpha}{K}\sum_i \frac{f_i}{c_i}\left[\log(\alpha/K) + \log(f_i/c_i)\right] \\
&= -(\alpha/K)\log(\alpha/K)\sum_i \frac{f_i}{c_i} - (\alpha/K)\sum_i \frac{f_i}{c_i}\log\frac{f_i}{c_i} \\
&= -\log(\alpha/K) - (\alpha/K)\sum_i \frac{f_i}{\langle c \rangle}\left(1 + \frac{\Delta c_i}{\langle c \rangle}\right)^{-1}\left[\log\frac{f_i}{\langle c \rangle} - \log\left(1 + \frac{\Delta c_i}{\langle c \rangle}\right)\right] \\
&\approx -\log(\alpha/K) - (\alpha/K)\sum_i \frac{f_i}{\langle c \rangle}\left(1 - \frac{\Delta c_i}{\langle c \rangle}\right)\left[\log\frac{f_i}{\langle c \rangle} - \frac{\Delta c_i}{\langle c \rangle}\right], \qquad \text{(S66)}
\end{aligned}$$

where we kept only the first power of $\Delta c_i/\langle c \rangle$. Rearranging the terms and still keeping only the first order terms, we can find

$$H|_{\delta=0} = H_F + \left[\left\langle \frac{\Delta c}{\langle c \rangle}\log f \right\rangle_f - \langle \log f \rangle_f \left\langle \frac{\Delta c}{\langle c \rangle} \right\rangle_f\right] + O\left((\Delta c_i/\langle c \rangle)^2\right), \qquad \text{(S67)}$$

which results in equation (15) from the main text.
